# Genome divergence between European anchovy ecotypes fuelled by structural variants originating from cross-hemispheric admixture

**DOI:** 10.1101/2024.09.15.613121

**Authors:** Laura Meyer, Pierre Barry, Alan Le Moan, Christine Arbiol, Rita Castilho, Carl Van der Lingen, Malika Chlaïda, Niall J. McKeown, Bruno Ernande, François Bonhomme, Pierre-Alexandre Gagnaire, Bruno Guinand

## Abstract

The formation of ecotypes is shaped by mechanisms that reduce gene flow through complex interactions between ecological, historical, and genomic factors. In the European anchovy (*Engraulis encrasicolus*), marine and coastal ecotypes have been identified in the North-East Atlantic and Mediterranean Sea, yet the genomic basis of their divergence remains unclear. Here, we present the first genome-scale analysis of this species complex, integrating whole-genome sequencing (WGS) and RAD-seq data from populations across its distribution range. In addition to the marine and coastal ecotypes, we identify a previously undetected lineage which is present in southern Morocco, the Canary Islands and even in South Africa. This southern Atlantic lineage exhibits a gradient of admixture with northern populations near the Atlantic-Mediterranean transition zone. Genomic differentiation landscapes reveal large regions of high linkage disequilibrium, likely corresponding to thirteen structural variants (SVs) segregating within or between the lineages. Notably, three of the six SVs contributing to the gene flow barrier between northern ecotypes originated in the southern lineage, supporting a partially shared evolutionary history between the coastal ecotype and the southern lineage. Our findings suggest that anchovy ecotype divergence has been shaped by a combination of ancient structural variation, admixture, and local adaptation. This study highlights how SVs that arose between geographically isolated lineages can act as key genetic elements in ecotype formation, reinforcing reproductive isolation through distinct evolutionary pathways.

## Introduction

Speciation is a complex, multi-level process shaped by temporal, spatial, ecological and genome architectural factors (Abbott et al., 2013). These dimensions collectively influence a variety of reproductive isolation mechanisms, which establish and accumulate barriers to gene flow, strengthening reproductive isolation during the course of speciation (Kulmuni et al., 2020; Schluter & Rieseberg, 2022). Although the multifaceted nature of speciation is widely recognized, many comprehensive studies have focused on systems where divergence appears to be primarily driven by one or a few dominant factors within a well-defined eco-evolutionary context. Ecotype formation is one such example, where genomic variation associated with distinct ecological habitats has traditionally been used to investigate ecologically driven speciation. In these cases, speciation is often assumed to be rapidly initiated by local adaptation, provided divergent selection arises from habitat variation and sufficient genetic variation is available for selection to act upon (Rundle & Nosil, 2005; Schluter, 2000; Schluter & Conte, 2009). However, recent advances in speciation genomics present a more nuanced perspective, highlighting growing evidence that ecotype formation results from the interplay of multiple factors (Van Belleghem et al., 2018; Marques et al., 2019).

A key finding from these genomic investigations relates to the temporal dimension of ecotype formation, which reveals that ecotype divergence is often associated with ancient allelic variation. Even when ecotype formation was initially attributed to the recent establishment of new habitats, divergence times are frequently found to exceed those derived from paleoenvironmental reconstructions or from coalescent time distributions for sequences from a panmictic population (Han et al., 2017; Le Moan et al., 2021; Louis et al., 2020; Nelson & Cresko, 2018; Small et al., 2023). This discrepancy between ecological and genetic divergence time frames (Van Belleghem et al., 2018) can be explained by several mechanisms, including long-term balancing selection on the variants underlying ecotype divergence, ancestral population structure or admixture between divergent lineages.

Another recurrent finding is that genetic differences contributing to ecotype formation are concentrated in genomic regions with low recombination rates, rather than being spread across the genome. These regions often comprise structural variants (SVs) such as chromosomal inversions, which appear to play a disproportionate role in ecotype divergence (Faria et al., 2019; Hager et al., 2022; Lundberg et al., 2023; Todesco et al., 2020). Recent evidence suggests that SVs may form a substantial barrier to gene flow, ensuring at least moderate reproductive isolation (RI) between lineages, particularly when multiple SVs separate the lineages and when these variants are coupled (Le Moan et al., 2024). Since recombination is largely suppressed in SVs such as inversions, divergence accumulates more readily between haplotypes, which are protected from the homogenizing effects of gene flow (Berdan et al., 2024; Rieseberg, 2001). While SVs are often implicated in RI between ecotypes through their role in local adaptation by capturing multiple loci under environmental selection (Kirkpatrick & Barton, 2006; Wellenreuther & Bernatchez, 2018), they may also harbor coadapted gene complexes or Dobzhansky-Muller incompatibilities that contribute to the isolation of nascent lineages carrying different haplotypes (Navarro & Barton, 2003).

Understanding the origin of the genetic variation that underlies ecotype divergence can be a first step towards reconciling the different components of ecotype formation. This can aid in identifying the most plausible evolutionary scenarios to explain how ecotypes have formed. One scenario that has been proposed to account for both the temporal and ecological components of ecotype formation, is admixture between geographical lineages. Admixture is not merely a homogenizing process; it has also been linked to evolutionary diversification that may lead to ecotype formation through several mechanisms. These include the transfer of locally adaptive variants via introgression (Abbott et al., 2013), combinatorial speciation through the emergence of new allelic combinations selected in different habitats (Marques et al., 2019), and the resolution of genomic conflicts caused by incompatibilities, resulting in RI from both parental lineages (Schumer et al., 2015). The latter may be followed by ecological niche segregation to reduce competitive interactions and avoid detrimental hybridisation. In such scenarios, SVs may provide “pre-packaged” divergent haplotypes that can readily contribute to RI following introgression (Edelman et al., 2019; Feder et al., 2003) or may better resist re-homogenization, compared to the collinear genome, after the onset of gene flow (Machado et al., 2007; Noor et al., 2001; Rafajlović et al., 2021). However, further empirical evidence is required to better understand the role of admixture in driving ecotype divergence through the transfer or assortment of SVs.

Here, we investigate the role of SVs in ecotype formation in the European anchovy, *Engraulis encrasicolus sensu lato*. This species complex has formerly been shown to be subdivided into a marine ecotype (offshore and pelagic, *E. engraulis s. stricto*) and a coastal ecotype (nearshore, lagoonal and estuarine, *E. maeoticus*) that are able to co-exist in quasi-sympatry despite frequent hybridisation (Le Moan et al., 2016). Partial RI between these ecotypes is evidenced by differences in both their genetic makeup and phenotypic characteristics (see Bonhomme et al., 2022 for a review). The divergence between ecotypes involves a condensed genomic architecture, suggesting a potential role for SVs in ecotype formation (Le Moan et al. 2016). Moreover, the most likely demographic model for their divergence involves secondary contact after a prolonged period of geographic isolation. The current distribution of *E. encrasicolus* spans the North-East Atlantic, Mediterranean, and Black seas. Classically, the southern range limit of *E. encrasicolus* along the western African coast is considered as the south of the Gulf of Guinea, while *E. capensis* has been described further south in the Benguela system off South Africa. However, some samples collected near the Atlantic-Mediterranean transition zone, off the Moroccan coast or near the Canary Islands, demonstrated genetic proximity with *E. capensis* (Silva et al., 2017; Zarraonaindia et al., 2012), suggesting that these might present a third unknown ancestry. These findings warrant further investigation into potential admixture scenarios between *E. capensis* and European anchovies, and considering possible evolutionary impacts on the divergence between the marine and coastal ecotypes.

We present the first study using whole-genome sequencing (WGS) in the *E. encrasicolus* species complex to provide a detailed characterization of its genetic structure. By employing reference-based mapping and anchoring to a chromosome-scale reference genome, we reveal the genomic architecture underlying divergence between the different anchovy lineages. To complement the WGS approach, we incorporated RAD-sequencing data to characterise the eco-geographic structure of anchovies across a broad geographical range. This included individuals from both the marine and coastal ecotypes in the northern part of the distribution, as well as anchovies from the Canary Islands, the Moroccan coast, and South Africa. We aimed to determine whether and how these lineages have genetically interacted over their evolutionary history and to evaluate the potential role of SVs in driving their divergence.

## Materials and methods

### Sampling and DNA extraction

Samples were collected from multiple sites covering a large part of the species distribution area (Locations table, **Supplementary Table S2**) and were issued from various sampling expeditions and local fisheries (**Supplementary Table S1**, Type==”Tissue”). These samples were collected in different types of habitats, which were classified either as coastal or marine habitats. Also included in our sampling scheme were eight individuals collected off the South African coast (Gqeberha). Whole genomic DNA was extracted from muscle tissue or fin clips using commercial tissue kits (Qiagen and Macherey-Nagel). Extraction quality was checked on agarose gel for the presence of high molecular weight DNA, and double-stranded nucleic acid concentration was measured using Qubit 2.0 and standardised in concentration before library construction.

### Reference genome assembly

We performed high-coverage linked-read (10X genomics) sequencing of a marine Atlantic *E. encrasicolus* individual from the Faro location (Algarve) to generate a new reference genome assembly (hereafter called Eencr_V1), following the same methodology as in Meyer et al. (2024). The *de novo* assembly obtained by analysing preprocessed linked-reads (raw coverage ∼40X) with supernova v2.1.1 (Weisenfeld et al., 2017) reached a total length of ∼926 Mb (925,873,119 bp; contig N50=13.08 kb; scaffold N50=20.36 kb). All downstream analyses required for variant calling from RAD-seq and WGS data were performed on the subset of scaffolds longer than 10 kb to account for assembly fragmentation. Genomic landscapes of differentiation and local PCA (see below) were reconstructed after anchoring scaffolds to the recently released chromosome-level assembly of an *E. encrasicolus* individual from the Black Sea (GenBank assembly accession: GCA_034702125.1) (**Supplementary Fig. S1**). Whole-genome alignment between our Eencr_V1 reference genome and the new assembly was performed with Minimap2 (Li, 2018) and visualised using D-GENIES (Cabanettes & Klopp, 2018).

### Whole-genome resequencing data

Thirty-nine samples (**Supplementary Table S1**, WGS==”yes”) were selected for whole-genome sequencing (WGS), including samples from coastal and marine habitats in the Atlantic (*GAS*) and the Mediterranean Sea (*GDL*, *SPN*). We also included samples from the Atlantic-Mediterranean transition zone (*PRS*) and from South Africa to investigate genetic makeup in these localities.

Individual WGS libraries were prepared following the Illumina TruSeq DNA PCR-Free Protocol and sequenced to an average depth of ∼10-30X on an Illumina NovaSeq 6000. Raw demultiplexed reads were processed using fastp (v0.19.05) (Chen et al., 2018) and aligned to our Eencr_V1 reference genome using BWA-MEM (BWA v0.7.17; Li, 2013). Picard (v2.26.8) (“Picard toolkit”, 2019) was used for sorting read alignments, marking duplicates and adding read groups.

Variants were called using the GATK best practices workflow (McKenna et al., 2010; Van der Auwera et al., 2013), without performing variant and base quality score recalibration steps. Firstly, individual GVCF files were created from bam files with HaplotypeCaller (GATK v.4.3.0.0). This information was then stored in a GVCF database using GenomicsDBImport, and VCF files (one file per scaffold) were generated with GenotypeGVCFs. These files were concatenated into a single VCF file which was filtered using vcftools v0.1.16 (Danecek et al., 2011) and bcftools v1.19 (Danecek et al., 2021) to retain only high quality SNPs. This included recoding genotypes as missing for low-quality calls (“--minGQ 20”) and hard-filtering sites based on: normalised variant quality (“QD<2.0”), mapping quality (“MQ<40.0” and “MQRankSum”), the rank sum test for site position within reads (“ReadPosRankSum<-8.0”), strand bias (“FS>60.0” and “SOR>3.0”), average genotype quality (“AVG(FMT/GQ)<20”), and average depth (greater than 90X, corresponding to the 97.5^th^ quantile of the individual read depth distribution, i.e. “AVG(FMT/DP)>90.0”). The VCF was finally filtered for indels (including nearby SNPs, “--SnpGap 5”), multiallelic SNPs (“--max-alleles 2”) and sites containing more than 15% of missing genotypes (“--max-missing 0.85”). The final VCF file (hereafter referred to as the **WGS dataset**) contained ∼5.9 M sites located on 9093 different scaffolds longer than 10 kb.

### RAD sequencing data

RAD-seq libraries were prepared for 243 samples (**Supplementary Table S1**, RAD==”yes” and Type==”Tissue”) by batches of 64 multiplexed individuals, following a similar protocol to Baird et al. (2008) using the *Sbf*I restriction enzyme. Twenty-five of these samples were also used to produce WGS data, providing a link to understand the genetic structure in both datasets. RAD library sequencing was performed on an Illumina HiSeq2500 sequencer in 100 bp single-read mode. To complement our sampling, we also included raw sequencing data for 128 anchovy samples from Le Moan et al. (2016), which were generated using the same restriction enzyme. The data were demultiplexed using process_radtags (Stacks v2.60) and reads were aligned to our Eencr_V1 reference genome using BWA-MEM (BWA v0.7.17; Li, 2013). The reference-based Stacks pipeline was run in an integrated workflow developed by the MBB bioinformatic platform (https://web.mbb.cnrs.fr/subwaw/workflowmanager.php) (Penaud et al., 2020). Gstacks was run using default parameters (“--model marukilow--var-alpha 0.05 --gt-alpha 0.05 --max-clipped 0.2”) with the minimum PHRED-scaled mapping quality set to 20 (“--min-mapq 20”). Thereafter, genotypes were exported in VCF format using the populations module (“--min-populations 2 --min-samples-per-pop 0.7 --min-maf 0.05”) and filtered to remove SNPs with more than 15% missing data across individuals. The VCF was also filtered to only retain sites that were present in the WGS dataset, since our objective was to describe the same genetic variation but at a larger geographic scale.

Lastly, all samples from the WGS dataset were integrated into the RAD VCF. The final VCF file (hereafter referred to as the **RAD dataset**) contained genotype data for 385 samples at 3880 variable sites.

### Population structure

To describe the genetic structure in both WGS and RAD datasets, we conducted genome-wide and chromosome-wide Principal Component Analysis (PCA) using the R package SNPRelate (v1.28.0) (Zheng et al., 2012). We used ADMIXTURE v1.3.0 (Alexander et al., 2009) to estimate individual ancestry proportions in all samples using the RAD dataset, assuming *K*=3 parental ancestries and using default parameters. Individual ancestry proportions were visualised in a triangle plot based on their ternary coordinates, and samples were then classified into different non-admixed and admixed ancestry categories based on their positions. The genetic composition of each sampling location was summarised by representing the fraction of individuals assigned to each ancestry category, in order to describe the eco-geographical distribution of the three ancestries.

Genetic differentiation (*F*_ST_), nucleotide diversity (*π*), absolute genetic divergence (*d*_XY_), and individual heterozygosity were calculated for the WGS data in non-overlapping 5 kb windows (with “—minSites 15”) using the popgenWindows.py script (Martin, 2018; https://github.com/simonhmartin/genomics_general).

### Identification and genotyping of structural variants

Our analyses of anchovy population structure revealed the presence of large SVs that were several megabases in size and that segregate across the distribution range. We therefore used chromosome-wide PCA to identify clusters of individuals representing alternate genotypes at each SV, and assigned individuals’ genotypes based on their cluster membership using both the WGS and RAD datasets. The PCA axis representing structural variation was evidenced by the presence of three separated clusters, and was in most cases supported by PC1. Samples which did not show clear cluster membership based on their PCA coordinates were not genotyped. For samples that had WGS data, we corroborated genotype assignment with the relative positions of each sample in local PCA, which was conducted in non-overlapping windows of 5 kb using lostruct (v0.0.0.9; Li & Ralph, 2019). In total, we genotyped individuals at 13 large SVs that occur on different chromosomes.

### Divergence history of structural variants

To study the evolutionary relationships between individuals carrying different haplotypes at the 13 SVs, we constructed neighbour-joining phylogenetic trees for each chromosome carrying a large SV. Trees were obtained using the “phylo” command from VCF-Kit (Cook & Andersen, 2017) which uses variable sites from a VCF to create a multiple sequence alignment to then calculate a difference matrix using MUSCLE (Edgar, 2004). Because our chromosome-wide PCAs indicated low recombination between alternate SV haplotypes on the 13 chromosomes, these SV trees can be used to resolve the evolutionary relationships among haplotypes without the confounding effect of recombination. For these analyses, we used a subset of high coverage individuals that are homozygous for the SV to avoid phasing issues and to facilitate visualisation of haplotype relationships. We rooted trees on the branch separating alternate homozygote groups. For comparison, trees were also constructed for chromosomes not carrying SVs and rooted using a South African sample (“ATL_MAR_ZDA_61_1162”).

## Results

### Genetic structure in the *E. encrasicolus* species complex reveals three-way admixture

We used a combination of both WGS and RAD-seq data as complementary approaches to study the genetic structure of anchovies in the eastern Atlantic, Mediterranean and Black Seas. Our RAD dataset, with mean per-sample coverage of 52.7X, revealed a clear picture of the overall genetic structure in the *E. encrasicolus* species complex across its entire range distribution. These results, obtained with a reduced representation SNP dataset, were very similar to those obtained using our WGS dataset (per-sample coverage 10-30X) which contains a smaller subset of samples (detailed results provided in the supplied HTML report, see **Supplementary Appendix**). Firstly, we observed genetic differentiation between samples collected in marine and coastal habitats in the northern part of the range, corresponding to the previously described marine and coastal ecotypes (Le Moan et al. 2016; Bonhomme et al. 2022). This can be observed along the second axis of variation (PCA 2 in **Fig. 1A**), where most coastal samples are positioned at the top of the plot, while marine samples fall in the bottom right corner. As for PCA 1, this axis shows a different signal that reflects geographic structure rather than ecological structure. On the horizontal axis, South African samples and other individuals collected off the African Atlantic coast (Morocco and the Canary Islands) are spread out towards the left-hand side of the plot, while the majority of other samples group to the right. Hence, PCA at a genome-wide scale shows the existence of three distinct genetic ancestries, which were further confirmed using admixture analysis. Inferred individual ancestry was represented in a ternary plot (**Fig. 1B**) showing the relative proportions of coastal (top), marine (right) and southern (left) ancestry for each individual. Ongoing gene flow between the three ancestries was revealed by considerable levels of admixture, in particular between the marine and coastal ancestries. A large number of samples also fell in the central area of the plot due to having balanced proportions of the three ancestry components, reflecting the existence of three-way admixture.

**Fig. 1.**
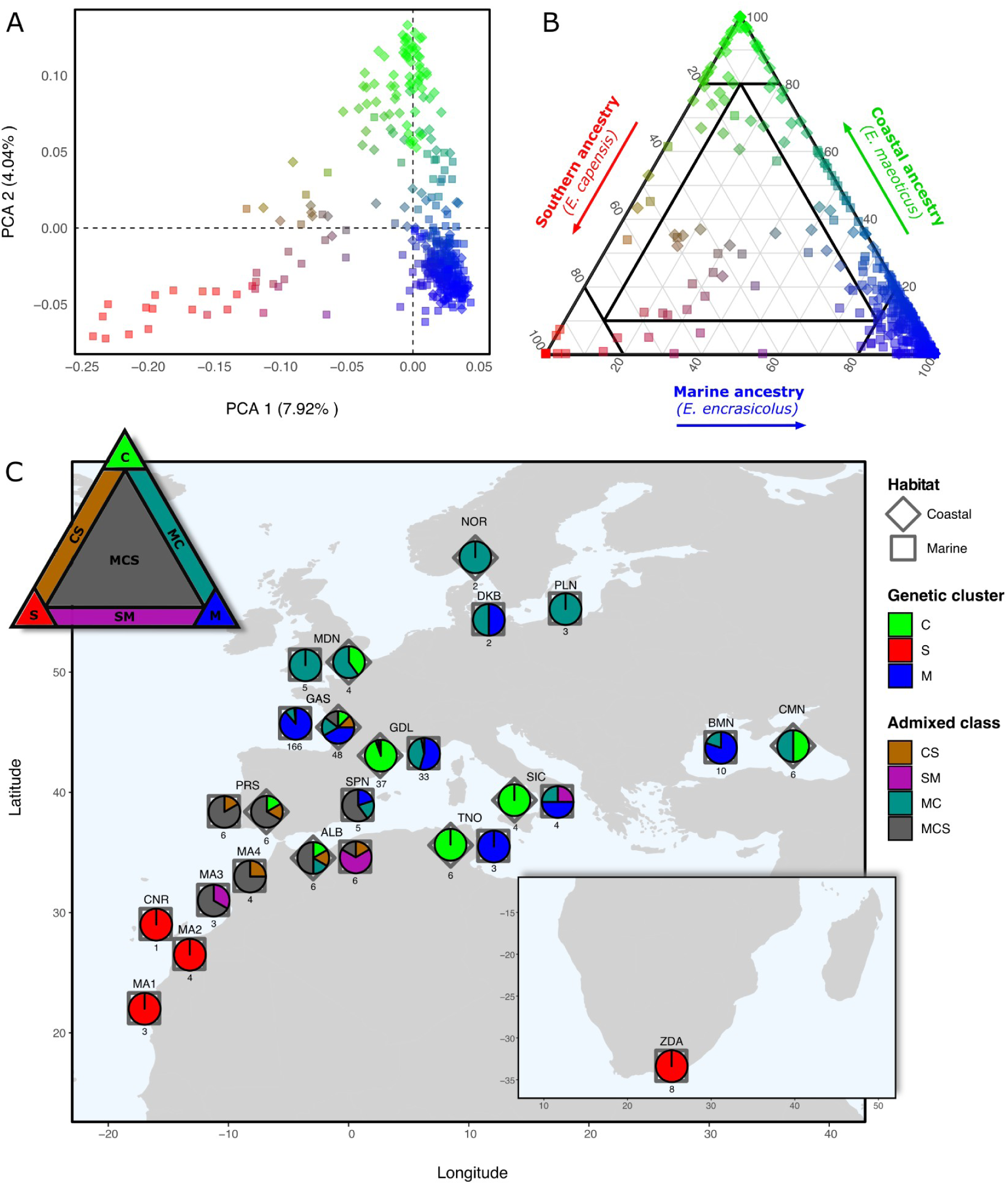
A) Principal Component Analysis (PCA) performed on the entire dataset of 385 anchovy samples. Sites used in the analysis were high-quality variants present both in the WGS data as well as in the RAD data, corresponding to a total of 3881 SNPs. Shapes indicate habitat type and colours reflect ancestry proportions as determined by admixture analysis (see B). B) Ternary plot showing the admixture level between three genetic ancestries: coastal (green), southern (red), and marine (blue) ancestry. Coordinates, as well as RGB colours, reflect the relative ancestry proportions of samples along each of the three axes. Samples were classified as belonging to a genetic cluster (black lines demarcating seven areas) based on their position in the plot. Clusters *C*, *S* and *M* represent “non-admixed” parental lineage ancestries, while *CS*, *SM*, *MC* and *MCS* represent various levels of admixture. B) Map with sampling locations where symbols represent habitat type and pie charts show the proportions of different genetic clusters present. Numbers beneath pie charts indicate sample sizes. Locations are described in **Supplementary Table S2**.

Based on their ternary coordinates, samples were classified as belonging to one of seven ancestry categories, each corresponding to a different sub-area of the triangle plot (black demarcations **Fig. 1B**, triangle in **Fig. 1C**). A sample was thus considered to belong to a non-admixed genetic cluster (coastal/*C*: green; southern/*S*: red; marine/*M*: blue) if that ancestry reached more than 80% of its total genetic ancestry. Secondly, we distinguished three different admixed classes where individual ancestry proportions were mainly made up of two ancestries (the third not amounting to more than 10%). These classes were *CS* (admixed between *C* and *S*; brown), *SM* (admixed between *S* and *M*; purple) and *MC* (admixed between *M* and *C*; seagreen). A last admixed class, called *MCS*, consisted of individuals with balanced proportions of all three ancestries (admixed between *M*, *C* and *S*; grey). The eco-geographical distribution of the three ancestries revealed that individuals belonging to the *C* cluster (green) were only found in coastal habitats (diamond symbols) in the northern part of the range, while *M* individuals (blue) mainly occurred in marine environments (square symbols) of the same region (**Fig. 1C**). This corresponds to the previously described ecotypic structure between coastal and marine anchovies. This structure was particularly evident in the Mediterranean Sea, where almost all coastal samples were part of the *C* cluster (e.g. coastal habitats in *SIC*, *TNO* and *GDL*). However, this signal of ecotypic differentiation becomes diluted near the Atlantic-Mediterranean boundary, where a gradient of increasing southern ancestry is observed. This admixture gradient can be seen through the increasing proportion of *MCS* individuals (grey) in the Alboran Sea (*ALB*), off the southern coast of Portugal (*PRS*) and in northern Morocco (MA4 and MA3). Finally, we observed that samples from locations to the south of the Canary islands (*CNR*), including the sampling site in South Africa (*ZDA*, inset map), all belonged to the *S* cluster (red).

After describing the three ancestries as well as their ecogeographic distribution patterns, we aimed to study their genomic architecture of differentiation. Genomic differentiation landscapes reconstructed between the coastal, marine and southern clusters using WGS data, yielded highly heterogeneous patterns that strongly varied from chromosome to chromosome (**Fig. 2**). The background level of differentiation between the marine and coastal clusters was lower (**Fig. 2A**, mean *F*_ST_ 0.013) compared to the background *F*_ST_ between the southern and coastal clusters (**Fig. 2B**, mean *F*_ST_ 0.075) and between the southern and marine clusters (**Fig. 2C**, mean *F*_ST_ 0.066) (see **Supplementary Fig. S9** for *F*_ST_ distributions). Genomic landscapes of differentiation between marine and coastal individuals in the Atlantic (**Fig. 2A**, *ATL*: first row of each comparison) and in the Mediterranean Sea (**Fig. 2A**, *MED*: second row) were similar, even though some differences were observed (e.g. on chromosomes CM068262 and CM068273).

**Fig. 2.**
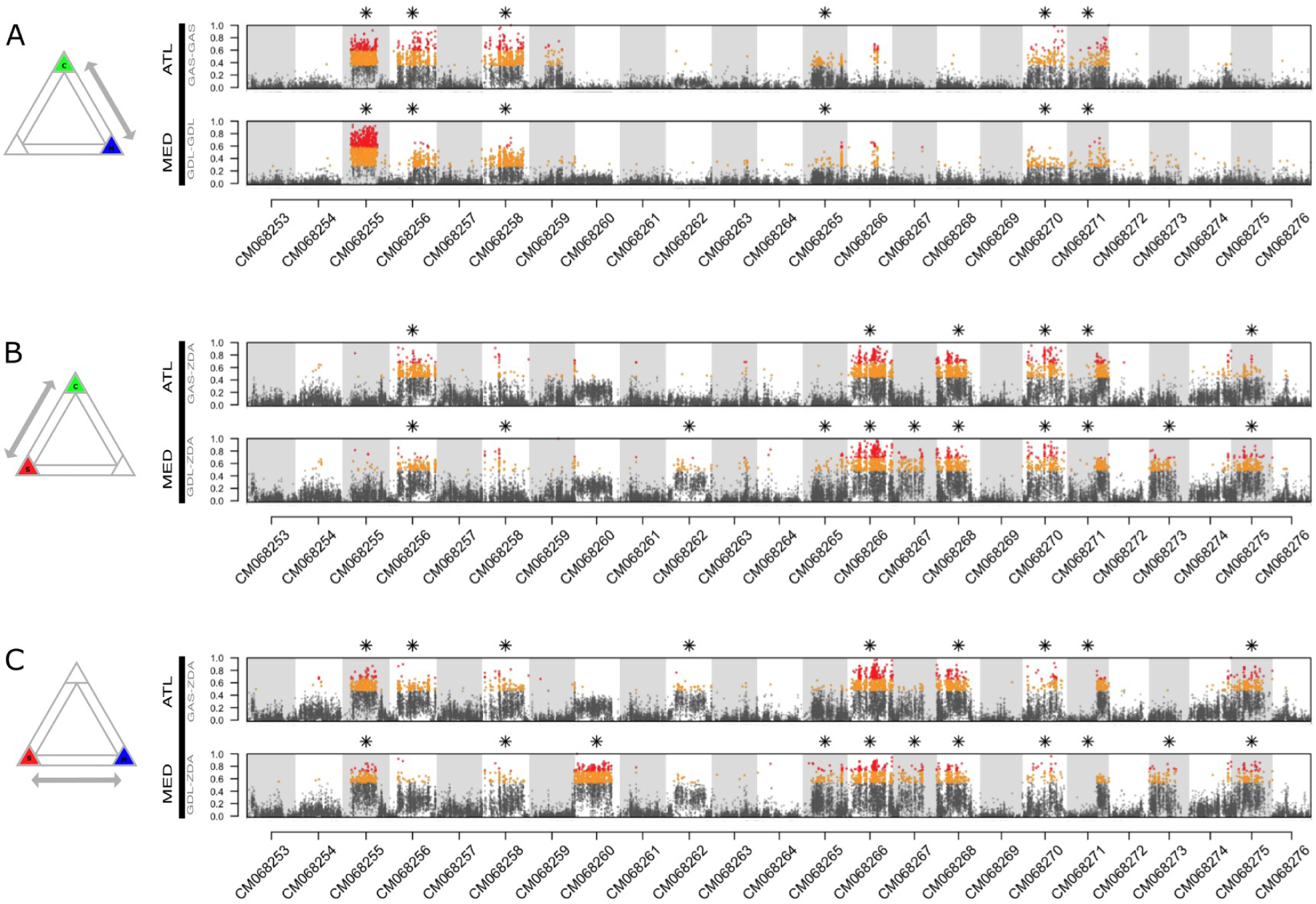
Genomic landscapes of differentiation (*F*ST) calculated in 5 kb sliding windows between groups of samples (3 individuals per group) from different genetic clusters (see Fig. 1). Differentiation landscapes are shown for three different comparisons (A: Coastal vs. Marine; B: Coastal vs. southern; C: southern vs. Marine). Each panel consists of two rows, representing cases where Coastal/Marine samples either originated from the Atlantic (ATL) or from the Mediterranean Sea (MED). Orange points are windows where *F*ST was higher than the 95^th^ quantile, while red points are above the 99^th^ quantile. Stars indicate chromosomes where more than 2.5% of windows showed *F*ST higher than the 95^th^ quantile. Grey and white rectangles delimit the 24 chromosomes of *E. encrasicolus*.

Heterogeneous *F*_ST_ values across genomic differentiation landscapes were largely characterised by the presence of high-differentiation regions (*F*_ST_ values above the 95^th^ quantile) which clustered into continuous *F*_ST_ plateaus. These patterns could point to the presence of large SVs that are associated with ecotype or lineage divergence. To assess whether these putative SVs harbour divergent, non-recombining haplotypes, we performed PCA on individual chromosomes using both the WGS and RAD datasets (**Supplementary Fig. S2 and S4**). Chromosome-wide PCAs revealed consistent patterns of clustering between the WGS and RAD datasets, but these varied greatly across chromosomes, as did the proportion of genetic variation explained by the first two PC axes.

Representative examples of chromosome-wide PCAs (**Fig. 3A-H**) illustrate the main diversity of observed patterns. While some chromosomes showed largely continuous ancestry gradients (**Fig. 3A & E**), others exhibited several discrete clusters where samples are tightly grouped (**Fig. 3B-D & F-H**). These tight PCA clusters indicate that a large number of SNPs are in strong linkage disequilibrium (LD), resulting in the segregation of a limited number of non-recombining haplotypes. Combined with elevated levels of heterozygosity in the intermediate clusters (**Fig. 3K, Supplementary Fig. S11**) and the continuous clustering of genotypes observed across numerous consecutive local PCA windows (**Fig. 3L**), these patterns strongly support the presence of multiple large SVs.

**Fig. 3.**
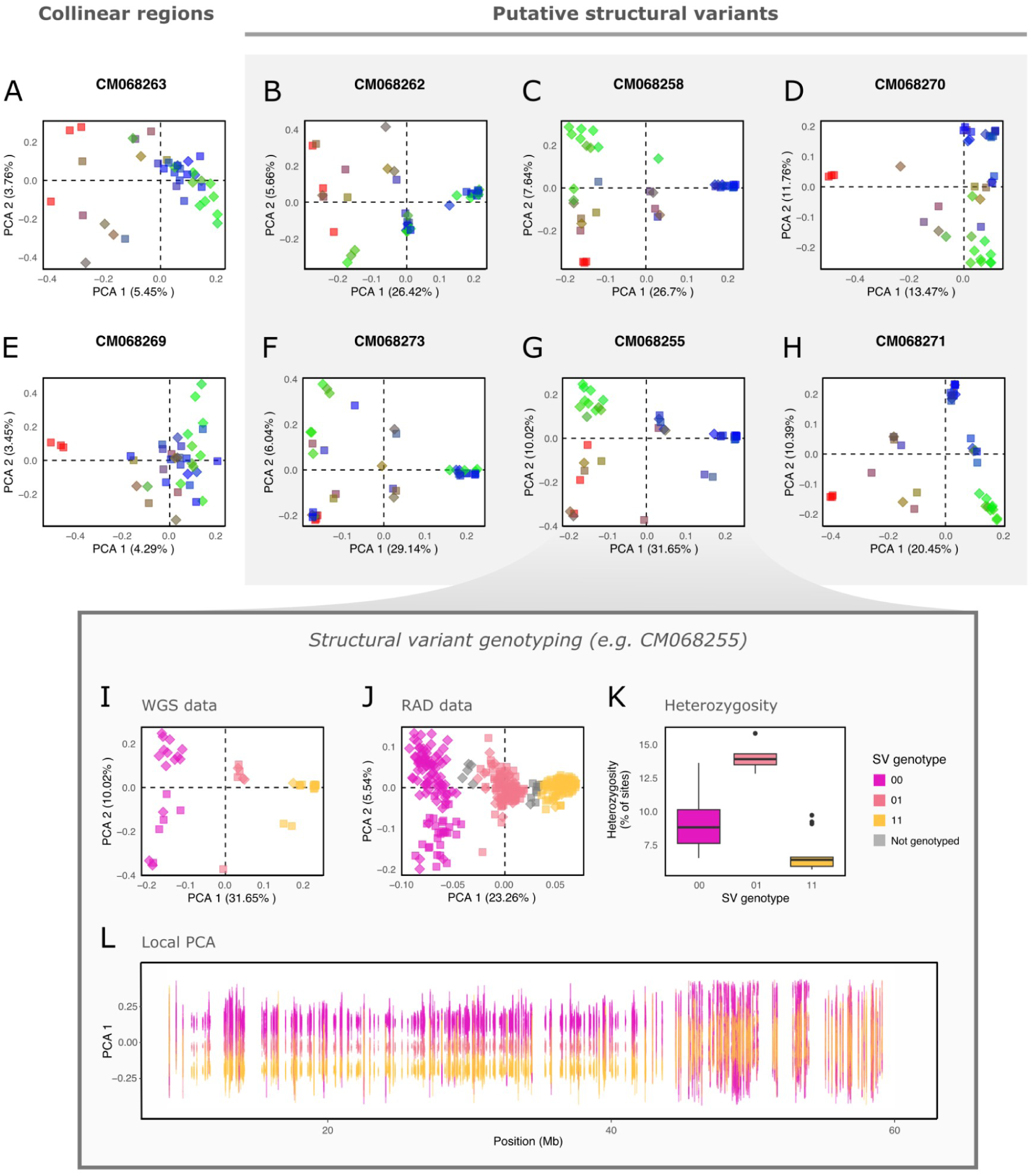
Examples of relatedness patterns on different chromosomes. PCA conducted on whole-genome data (n=39) at a chromosome-wide level (A-H) reveals that certain chromosomes show signs of reduced recombination through the presence of tight PCA clusters (B-D, F-H). These chromosomes show evidence for the presence of structural variants (SVs), which we aimed to genotype in all individuals (I-L). SV genotypes were assigned for whole-genome and RAD data (n=385) based on PCA coordinates (I & J). For whole-genome data, these assignments were further corroborated by individual heterozygosity levels (K) and local PCA patterns (L). In (L), PCA 1 coordinates plotted for non-overlapping 5 kb windows along the chromosome show sustained clustering patterns in the SV region (ending around ∼40 Mb). Gaps represent regions on CM068255 for which we did not have data (scaffolds < 10 kb or not present in our reference genome). Colours either reflect genome-wide ancestry proportions (A-H) or the SV genotype that was assigned to each individual (I-L).

In total, we identified 13 chromosomes (indicated by asterisks in **Fig. 2**) that showed evidence for SVs spanning at least 2.5% of the windows on the chromosome. Using individual coordinates from chromosome-wide PCAs, these 13 SVs were successfully genotyped (**Supplementary Fig. S3 and S5**) and individuals were classified as either *00* homokaryotes (pink), *01* heterokaryotes (salmon) or *11* homokaryotes (gold) (**Fig. 3I-L**). Individuals that could not be confidently assigned to any given group were not genotyped (grey). For consistency, we always assigned the *00* genotype to the group containing the most southern samples, in order to polarise the *0* haplotype with respect to southern lineage ancestry. For the SV located on chromosome CM068256, the PCAs based on the WGS and RAD datasets showed different results, which complicated our assignment of genotypes for this SV (**Supplementary Fig. S4 and S6**). We opted to genotype this SV according to the variation captured by PC1 in the WGS analysis, since this dataset included more markers and the structure was consistent with the phylogenetic reconstruction (**Supplementary Fig. S7)**.

### Anchovy lineages are differentiated at multiple SVs

Based on the assigned SV genotypes, we analysed the frequency patterns of the two haplotypes at each SV (*0* and *1*) (**Supplementary Fig. S8**) as well as the genotype frequencies (*00*, *01* or *11*) within each genetic cluster (**Fig. 4**). We observed that the coastal, marine and southern clusters (background colours in **Fig. 4**) carried different sets of SV genotypes. The southern cluster (bottom row of pie charts) largely harboured homokaryotic *00* genotypes at SVs (mean *00* frequency of 77% across SVs, 6 out of 13 fixed), while marine individuals (top two rows) were mostly homokaryotic for the *11* genotype (mean *11* frequency of 79% across SVs). In coastal individuals, on the other hand, some SVs were nearly fixed for the *00* genotype, while others were fixed for the *11* genotype (see details below). Overall, we found that the three clusters show substantial frequency differences at multiple SVs (**Supplementary Table S3**), which is coherent with the *F*_ST_ plateaus that were observed in the differentiation landscapes (**Fig. 2**). Admixed individuals that carry more than one type of ancestry (e.g. *MCS*) were often heterokaryotes at SVs (**Supplementary Fig. S6**), consistent with their admixed status.

**Fig 4.**
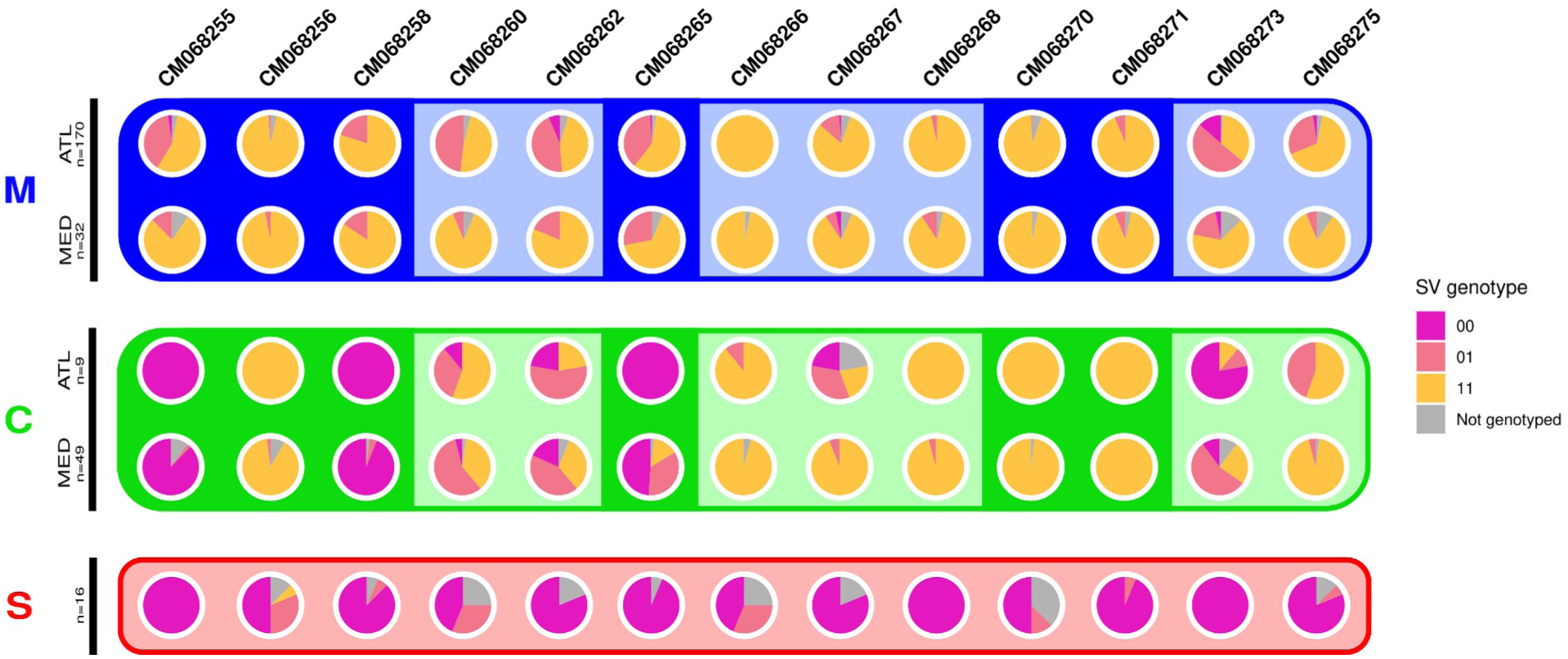
Genotype frequencies for 13 SVs on different chromosomes (columns). Pie charts show frequencies for the *00* (pink), *01* (salmon) and *11* (gold) genotypes in the Marine cluster (blue background), Coastal cluster (green background) and southern cluster (red background), with upper and lower rows corresponding to Atlantic and Mediterranean samples respectively (for *M* and *C*). Darker background colour in *M* and *C* indicates chromosomes that show elevated *F*ST when comparing Marine and Coastal individuals (Fig. 2A). For three of these chromosomes (CM068255, CM068258 and CM068265), ecotype differentiation involves southern haplotypes (*00*) that are present at high frequency in the Coastal individuals, while this is not the case for CM068270 and CM068271.

Even though SV frequency differences between the coastal, marine and southern clusters are observed, these SVs are not always fixed for a given haplotype but display a degree of haplotype sharing between the three clusters. This is clearly visible on CM068262 and CM068273, for example, where the SVs are polymorphic in almost all populations (**Fig 4**). Patterns of haplotype sharing can also be observed in neighbour-joining trees of these SV regions (**Fig. 5B & F**), where samples group according to their SV genotype (pink and gold branches) and not according to their ancestry category (tip colour). By looking at the general patterns of haplotype distributions across different SVs, it can be observed that southern ancestry haplotypes (*0*) are common in marine and/or coastal populations in the north. Furthermore, these southern haplotypes are slightly more common in the Atlantic (first and third rows in **Fig. 4**) than in the Mediterranean (second and fourth rows).

**Fig. 5.**
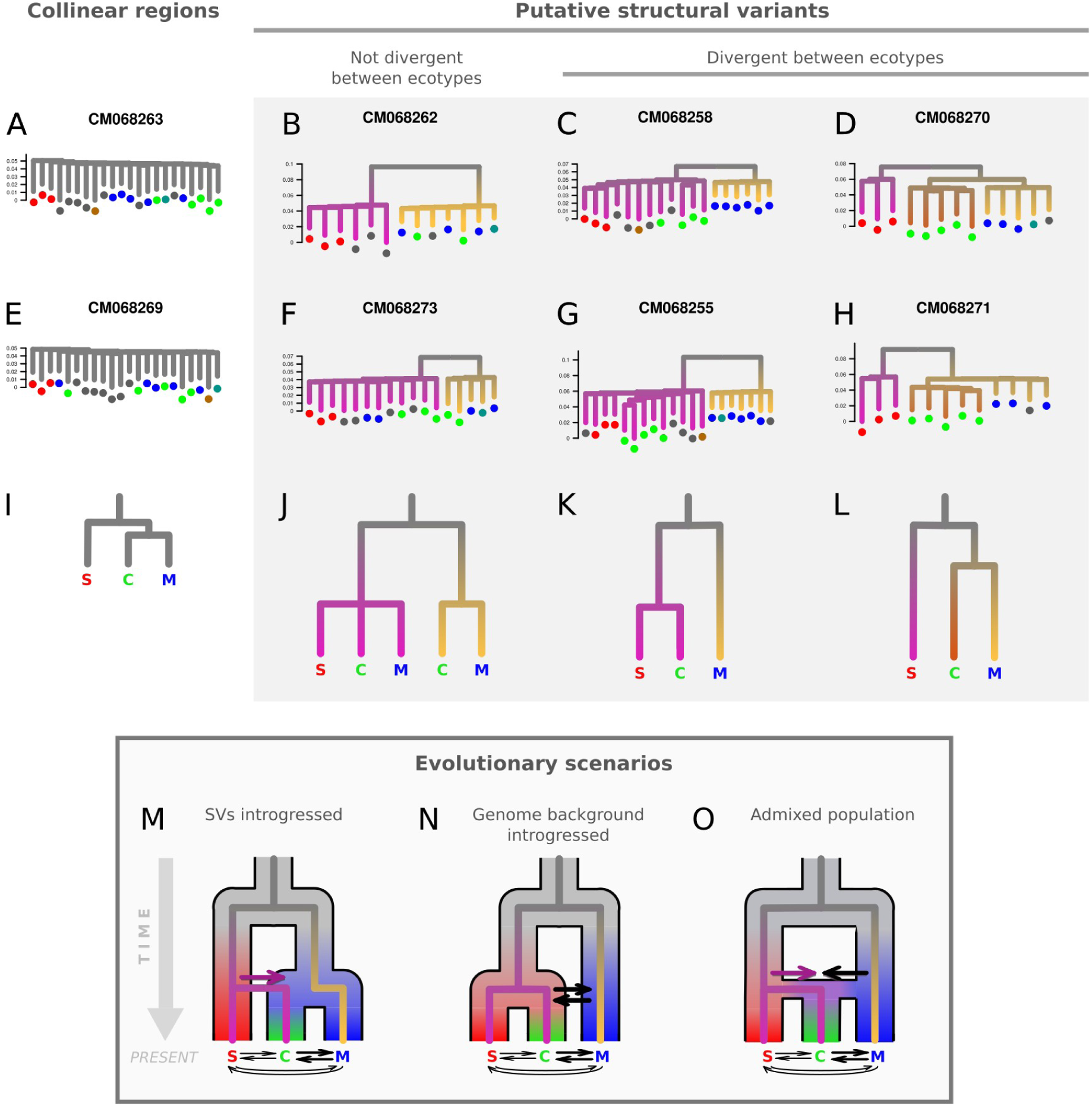
A-H) Neighbour-joining trees showing interindividual relationships on the same subset of chromosomes as in Fig. 3. Short branch lengths for normally recombining chromosomes (A & E) contrast with long branches between divergent haplotypes at SVs (B-D, F-H). These SVs show varying patterns of being shared or being private to the coastal, marine or southern clusters. This is what we illustrate with schematic trees in (I-L), where pink or gold colouration reflects divergence between haplotypes. In (L), brown colouration indicates the divergence of a third haplotype differentiating coastal and marine samples (e.g. CM068270 and CM068271). We propose three evolutionary scenarios (M, N & O) involving gene flow amongst lineages (horizontal arrows) which could explain the observed patterns of haplotype distributions across SVs. For (A & E), trees were constructed using SNPs on the entire chromosome, whereas in (B-D, F-H), the region was limited to that covered by the SV. For trees of SV regions, intermediate samples that present heterokaryotes are not displayed. Leaf tips are coloured according to individual ancestry categories. Trees were plotted on the same vertical scale.

One of our main findings is that there is a substantial excess of haplotype sharing at SVs between the southern and coastal clusters (mean *F*_ST_ 0.218 in SV regions), which is not observed between the southern and marine clusters (mean *F*_ST_ 0.273 in SV regions). This is particularly evident on CM068255, CM068258 and CM068265, where *0* haplotypes are highly predominant (sometimes fixed) in the coastal samples. These three chromosomes are among the six showing high *F*_ST_ between the marine and coastal clusters (asterisks in **Fig. 2A**), which can thus be explained by the presence of southern haplotypes in coastal samples. This was confirmed in phylogenies of the SV regions (**Fig. 5C & G**), since coastal and southern samples are grouped in the same branch (pink), while marine samples (with the alternative haplotype) are in a separate branch (gold).

On the other hand, high *F*_ST_ values between coastal and marine samples on CM068256, CM068270 and CM068271 are not due to the presence of southern haplotypes. On these chromosomes, northern populations almost exclusively carried *1* haplotypes (rows 1-4 in **Fig. 4**) that were, in addition to their differentiation from the *0* haplotype, also divergent between *C* and *M*. The presence of haplotype structure within the *1* haplogroup is indeed suggested by the separation of coastal and marine samples on PCA 2 (**Fig. 3D & H**), and also supported by the subdivision between different *1* haplotypes in the phylogenies (brown and gold branches in **Fig. 5D & H**). Certain SVs on other chromosomes also showed evidence for three distinct haplotypes (e.g. CM068273), but only CM068256, CM068270 and CM068271 were investigated (haplotype frequencies shown in **Supplementary Fig. S8)** since these chromosomes were involved in ecotype differentiation (**Fig. 2A**). Overall, these patterns suggest that there are three distinct haplotypes segregating on CM068256, CM068270 and CM068271, potentially resulting from the presence of multiple SVs on each chromosome.

By studying the distribution of *d*_XY_ values (**Supplementary Fig. S10**) and the branch lengths separating different haplotypes in phylogenies (**Supplementary Fig. S7)**, we gathered information about divergence in different genomic regions. We aimed to evaluate their concordance with alternative divergence scenarios and observed different types of patterns that are illustrated by the four columns in **Fig. 5**. We found that chromosomes that do not carry large SVs are characterised by short branch lengths and low levels of divergence (**Fig. 5A & E**). Such collinear regions did not show pronounced genetic structure between the three clusters and supported highest genetic similarity between the coastal and marine clusters. These patterns contrasted with phylogenies reconstructed from SV regions, where long branches were found to separate samples carrying different genotypes (**Fig. 5B-D & F-H**). Estimated *d*_XY_ values between alternate homokaryotes showed relatively similar distributions across SVs, with mean *d*_XY_ between 0.15 and 0.20%. Divergence between coastal *1* haplotypes and marine *1* haplotypes on CM068256, CM068270 and CM068271 was lower, as is reflected by shorter branch lengths in phylogenies (**Fig. 5D & H**). We note, however, that SV block delimitation (based on *F*_ST_ and local PCA patterns) was probably not precise enough to exclude all recombinant regions and to provide precise divergence estimates.

## Discussion

We present the first genome-scale study investigating genetic structure in the *E. encrasicolus* species complex, comprising anchovies from the eastern Atlantic, Mediterranean and Black Seas. Our findings reveal that two previously described ecotype lineages (marine and coastal) genetically interact with a third, southern lineage. This lineage has not previously been described as such, even though some anchovy samples from the northeast Atlantic and from southern Morocco were shown to display genetic proximity with *E. capensis* (Silva et al., 2017; Zarraonaindia et al., 2012). Although taxonomic considerations are beyond the scope of this study, our results show that the coastal, marine and southern lineages are primarily differentiated by genotype combinations at multiple large SVs, with the collinear parts of their genomes being weakly differentiated. We further found that these SVs most likely originated from admixture with the southern lineage, and that the coastal ecotype carries southern haplotypes at several SVs. Our study supports that SVs that arose in geographically isolated lineages can play an important role in driving ecotype formation following lineage admixture.

Previous genetic studies have shown that the European anchovy is subdivided into marine and coastal ecotypes that are present from the Bay of Biscay, through the Mediterranean to the Black Sea (reviewed in Bonhomme et al., 2022). Here, we show that there is a third component of genetic structure in this species complex, corresponding to an Atlantic lineage occurring off the African coastline. This southern lineage shows genetic homogeneity at a very large spatial scale, with genetic similarity between individuals sampled in Morocco, the Canary Islands and even as far as South Africa. However, from northern Morocco and southern Portugal into the Alboran Sea, we observe genetic admixture resulting in a gradient of decreasing southern ancestry. Previous studies reporting various signals of spatial structure in this species may have unknowingly captured different aspects of these complex admixed ancestries, leading to many different and conflicting interpretations in the literature. For instance, Zarraonaindia et al. (2012) reported the presence of an admixture gradient extending from the Canary islands to the northern Iberian peninsula. However, this gradient was attributed to the interaction of two components corresponding to populations inhabiting narrow versus wide shelf waters. We instead propose that this region corresponds to a three-way contact zone between the southern lineage and the two northern marine and coastal ecotype lineages, thus involving both an ecological component as well as admixture between geographic lineages. We observe post-F1 introgressive hybridisation resulting in widespread admixture and gene flow between the southern, coastal and marine genetic clusters, as is reflected by gradual ancestry gradients in the PCA plot (**Fig. 1**). Almost all individuals that were identified as belonging to the three-way admixed class (MCS) were sampled in the contact zone between the three lineages (**Fig. 1C**). This pattern of three-way admixture also extends further north in the Atlantic, where a few individuals carrying a MCS background were detected. The existence of gene flow between coastal and marine ecotypes has already been illustrated in previous work (Le Moan et al., 2016), but our results reveal that admixture with the southern lineage also contributes to global diversity patterns.

We found evidence for multiple megabase-scale SVs that segregate in the marine, coastal and southern anchovy lineages. Although the presence of SVs was only supported through indirect evidence based on LD, divergence and heterozygosity patterns, these regions showed many of the signals typically associated with chromosomal inversions (Mérot et al., 2020). In the literature, it has often been reported that SVs play a role in differentiating evolutionary lineages within species complexes, suggesting that they can play an important role in the formation of cryptic lineages or ecotypes (Berdan et al., 2024; Zhang et al., 2021). Our results in anchovies seem to point in this direction, since markers differentiating lineages and ecotypes were largely concentrated in SVs, whereas collinear regions of the genome showed low differentiation levels which likely reflects the homogenising effect of gene flow and recombination. By reconstructing the genomic landscape of ecotype divergence, we determined that marine and coastal ecotypes were differentiated at six large SVs. These SVs cover roughly 25% of the genome, which is in line with a previous study that estimated that the barriers to gene flow between ecotypes span 20 to 25% of the genome (Le Moan et al., 2016). The fine-scale structure of the SVs will need to be addressed by long-read sequencing in order to directly identify breakpoints and to resolve the possibly complex variation at each SV. In addition, there is also the possibility that small genomic islands of differentiation, located in the collinear genome and not detected here, could also play a role in ecotype divergence.

Our results also revealed that the origin of the six SVs that differentiate ecotypes is closely linked to the impacts of admixture between divergent geographic lineages. In particular, the coastal ecotype carries the same haplotype as the southern lineage at a minimum of three SVs. If these SVs were already segregating in the ancestral population to the marine, coastal and southern lineages, different conflicting genealogies would have been expected due to incomplete lineage sorting (ILS). However, we do not observe any SVs where the marine and southern lineages carry the same haplotype and are divergent with the coastal ecotype. If these patterns are not due to ILS, it suggests that the coastal and southern lineages could have a partly shared evolutionary history. Overall, the patterns of population structure in the collinear genome as well as haplotype distributions at SVs (whether shared between lineages or private) suggest that there are three alternative scenarios which could underlie the observed genealogical patterns (**Fig. 5M, N & O**):

1) In the first scenario, SVs could have been introgressed from the southern lineage into the coastal lineage. The deepest split in the tree represents the initial divergence between the southern lineage and the northern ancestral branch giving rise to the marine and coastal ecotypes (**Fig. 5M**). Following this split, genomic rearrangements likely accumulated in each branch, and subsequent gene flow could have led to the introgression of SVs. In this scenario, haplotypes from the southern lineage may have been introgressed into a preexisting coastal lineage or may have contributed to the formation of the coastal ecotype. This introgression likely occurred during an earlier period of contact, since current admixture in the contact zone seems insufficient to explain why coastal samples from across the distribution range almost exclusively carry southern haplotypes at these SVs. Evidence for the introgression of SVs is documented in other systems (Jay et al., 2018; Todesco et al., 2020), where they may serve as a source of divergent haplotypes involved in the evolution of novel traits or ecotype formation.
2) The second scenario proposes that the coastal ecotype directly originated from the southern lineage and does not share its most recent common ancestor with the marine lineage (**Fig. 5N**). In this case, recent common ancestry between the coastal and southern lineages would only be observed at the SVs, as widespread introgression with the marine lineage has eroded divergence in the collinear genome. This scenario does not require multiple SVs to successfully pass through various selective filters to establish. Instead, it involves neutral introgression in the background genome, as well as polymorphism at certain SVs that were shared via gene flow.
3) Another alternative scenario is whole-genome lineage admixture, where the coastal ecotype would have originated from an admixed population composed of both southern and northern ancestries. This admixture could have been followed by selective reassortment of SVs, driven either by adaptation to the coastal environment (Marques et al., 2019) or by the resolution of genomic conflicts resulting from incompatibilities (Schumer et al., 2015).

The evolutionary history of the marine and coastal ecotypes has previously been studied, and demographic inference has indicated that ecotype divergence probably occurred in a context of allopatric isolation followed by secondary contact (Le Moan et al., 2016). Although the aforementioned study did not include the third anchovy lineage in their analyses, the period of allopatric divergence that was inferred between ecotypes may in fact correspond to the split between northern and southern lineages (**Fig. 5M & N & O**). This divergence signal may have been preserved within the SVs, as is suggested by their *d*_XY_ distributions and long internal branch lengths in genealogies. By contrast, the signal of past divergence between the northern and southern lineages may have been eroded by subsequent gene flow and recombination in the background genome.

In addition to the six SVs that separate coastal and marine anchovies, we also identified other SVs (e.g. on CM068260) that are shared between ecotypes and where one haplotype has a southern lineage origin. Here again, the lack of recombination between rearrangements has preserved the historical divergence signal, despite the fact that these SVs have been subsequently exchanged between ecotypes. This result closely mirrors what has been described by studies using mitochondrial data, where two deeply divergent clades (called A and B) were found to segregate in populations now known to correspond to the marine and coastal ecotypes (Magoulas et al., 1996; Oueslati et al., 2014) and the southern lineage (Chahdi Ouazzani, 2016; W. S. Grant et al., 2005; Silva et al., 2014, 2017). Therefore, the mitochondrial genome and the shared SVs likely represent genomic regions that have retained the signal of north-south divergence despite gene flow following secondary contact.

The last type of genealogical pattern that we identified for certain SVs, separates the southern lineage from coastal and marine ecotypes that share more recent common ancestry (**Fig. 5D & H**), thus displaying the same topology as for the collinear genome but with significantly longer branches. This might imply that, in addition to north-south divergence, there is also more recent divergence between the two northern anchovy ecotypes, possibly involving complex rearrangements such as nested inversions (Joron et al., 2011; Kollar et al., 2024). Alternatively, these SVs could have a similar evolutionary history as proposed above in scenarios 1, 2 & 3. Namely, coastal anchovy would have carried a southern haplotype in the past, before it recombined with the northern haplotype due to gene flow between coastal and marine ecotypes, consequently decreasing the level of divergence between haplotypes in these populations. Gene flow between inversion arrangements has previously been described in several other systems, in particular for large chromosomal inversions that are impacted by gene conversion and double crossovers (Korunes & Noor, 2019; Matschiner et al., 2022; Meyer et al., 2024; Navarro et al., 1997).

Overall, we found that secondary contact between the northern and southern anchovy lineages ‒ whether involving introgression of specific SVs (scenario 1), background genome introgression of the southern lineage (scenario 2), or genome-wide admixture (3) ‒ is sufficient to explain the diversity of genealogical patterns observed at the genome-wide scale. However, our results do not allow us to determine which of the three alternative evolutionary scenarios is the most likely. This would require more in-depth demographic divergence analyses and complementary information regarding the past geographic distributions of the different lineages, especially during the last glacial period. However, we may hypothesise that the divergence between the northern and southern branches probably took place across hemispheres, as it broadly reflects the current distribution of the extant lineages.

Alternatively, divergence may have occurred between the northeast and -west Atlantic, since anchovy populations from the Gulf of Mexico may also form part of the *E. encrasicolus* species complex (Silva et al. 2014; 2017). It is likely that several periods of secondary contact could have taken place between ancient anchovy lineages, most likely during cooler periods when long-range distributional range shifts were possible and when the lineages were confined to lower latitudes due to ice sheets near the poles (Bouchenak-Khelladi et al., 2008; W. Grant & Bowen, 1998). In particular, the population of southern anchovies off the Moroccan coast could have originated during the last glacial episode, when anchovies from the southern hemisphere could migrate north and come into contact with northern populations.

It remains unresolved, however, whether the ancestral anchovy lineages were already associated with particular habitats or how/when this association took place to result in the current coastal and marine ecotypes. Today, the southern lineage seems to be associated with marine habitats, at least along the Moroccan, Mauritanian, Namibian and South African coasts (Chahdi Ouazzani et al., 2017; Whitfield, 1994). In this light, it is not obvious at first sight how the southern SVs haplotypes, which are carried by the coastal ecotype, could underlie adaptation to coastal environments. However, it is possible that epistatic interactions with loci in the genomic background have affected ecological traits that were not initially associated with these haplotypes. A classical example is the introgression of the Denisovan *Epas1* variant involved in adaptation to high altitude in modern humans, without being obviously involved in adaptation to altitude in the Denisova lineage (Huerta-Sánchez et al., 2014).

Alternatively, mutations could have accumulated in the SVs over time, conferring local adaptation to the coastal environment, which was not initially the case in the southern lineage. Empirical studies (Lamichhaney et al., 2016) and recent simulation work (Schaal et al., 2022) have indeed found support for the gradual accumulation of local adaptation alleles in inversions under high gene flow conditions.

Our study highlights the crucial role of SVs and admixture in the formation of ecotypes, supporting that hybridisation between geographically isolated lineages can provide the genetic substrate necessary for ecological specialisation and partial RI. SVs can act as “pre-packaged” divergent genomic elements containing multiple genes that contribute to reproductive barriers through local adaptation, coadapted gene complexes, or genetic incompatibilities. This supports a broader view that ecotype formation is rarely driven by strict *in situ* adaptation alone, but often involves phases of allopatric divergence and other historical contingencies (Van Belleghem et al., 2018; Bernatchez et al., 1996; Bierne et al., 2011; Hendry, 2009; Rundell & Price, 2009). Following lineage admixture, SVs can fuel ecotypic differentiation and resist gene flow over the long term, facilitating coupling among different components of RI. These associations are necessary for the maintenance of ecotypes and are instrumental if speciation between incipient ecotypes is ever to complete (Barton & de Cara, 2009; Bierne et al., 2011; Butlin & Smadja, 2018). Furthermore, the role of coupled SVs may be even more significant for the persistence of differentiated ecotypes in high gene flow marine species (Han et al., 2020; Jones et al., 2012; Kirubakaran et al., 2016; Wilder et al., 2020). However, whether these SVs alone are sufficient for marine ecotypes to speciate remains uncertain (Johannesson et al., 2024), as the moving balance between gene flow and divergent selection continuously reshapes their evolutionary trajectory. Future research should explore the broader role of lineage admixture as a source of divergent SVs and investigate the evolutionary history of these variants, to reveal how they contribute to the formation and maintenance of ecotypes across different taxa.

## Acknowledgements

We wish to thank the numerous colleagues that have provided us with samples and insightful contributions over the many years of this long-term study, more particularly amongst them Jean-Pierre Quignard, Karima Fadhlaoui-Zid, Petr Strelkov, Laurent Soulier, Lilia Bahri-Sfar and Nicolas Bierne. This work was supported by the ANR grant CoGeDiv ANR-17-CE02-0 006-01 to Pierre-Alexandre Gagnaire. We would like to thank the MGX-Montpellier GenomiX platform as well as the Montpellier Bioinformatics Biodiversity platform (MBB) supported by the LabEx CeMEB.

## Notes

### Competing Interest Statement

The authors have declared no competing interest.

### Summary of Updates

The main changes relate to how the introduction and discussion were framed (now we focus more on the role of admixture), the addition of a third possible evolutionary scenario (past existence of a completely admixed population), and more details were added in the results and supplementary figures.

