## Supplementary material for "Genome divergence between European anchovy ecotypes fuelled by structural variants originating from cross-hemispheric admixture"

### Supplementary information

#### Supplementary Tables

**Supplementary Table S1: Sample Table.** All samples used in the study (n=385), including 128 samples obtained from Le Moan et al. (2016). Habitat type was classified as either coastal ("COT", i.e. lagoons and estuaries) or marine ("MAR"). The "WGS" and "RAD" columns indicate whether a sample was included for whole-genome sequencing and/or RAD-sequencing. The last column corresponds to the genetic cluster or admixed class that each sample was assigned to based on ADMIXTURE results (**Fig. 1B**).

| NOM_FINAL | Type | Latitude | Longitude | Location | Habitat | WGS | RAD | Ancestry category |
|---|---|---|---|---|---|---|---|---|
| MED_MAR_ALB_34_0483 | Tissue | 35.256 | -3.835 | ALB | MAR | no | yes | SM |
| MED_MAR_ALB_34_0493 | Tissue | 35.256 | -3.835 | ALB | MAR | no | yes | SM |
| MED_MAR_ALB_34_0494 | Tissue | 35.256 | -3.835 | ALB | MAR | no | yes | SM |
| MED_LAG_ALB_37_1436 | Tissue | 35.170 | -2.870 | ALB | COT | no | yes | MCS |
| MED_LAG_ALB_38_1439 | Tissue | 35.170 | -2.870 | ALB | COT | no | yes | CS |
| MED_LAG_ALB_37_1431 | Tissue | 35.170 | -2.870 | ALB | COT | no | yes | MC |
| MED_LAG_ALB_37_1432 | Tissue | 35.170 | -2.870 | ALB | COT | no | yes | MCS |
| MED_LAG_ALB_38_1440 | Tissue | 35.170 | -2.870 | ALB | COT | no | yes | MCS |
| MED_LAG_ALB_38_1441 | Tissue | 35.170 | -2.870 | ALB | COT | no | yes | C |
| MED_MAR_ALB_35_0571 | Tissue | 35.313 | -2.661 | ALB | MAR | no | yes | MCS |
| MED_MAR_ALB_35_0584 | Tissue | 35.313 | -2.661 | ALB | MAR | no | yes | CS |
| MED_MAR_ALB_35_0585 | Tissue | 35.313 | -2.661 | ALB | MAR | no | yes | SM |
| BLS_MAR_BMN_39_0789 | Tissue | 43.701 | 29.436 | BMN | MAR | no | yes | M |
| BLS_MAR_BMN_39_0790 | Tissue | 43.701 | 29.436 | BMN | MAR | no | yes | M |
| BLS_MAR_BMN_39_0791 | Tissue | 43.701 | 29.436 | BMN | MAR | no | yes | MC |
| BLS_MAR_BMN_39_0792 | Tissue | 43.701 | 29.436 | BMN | MAR | no | yes | M |
| BLS_MAR_BMN_39_0793 | Tissue | 43.701 | 29.436 | BMN | MAR | no | yes | M |
| BLS_MAR_BMN_39_0794 | Tissue | 43.701 | 29.436 | BMN | MAR | no | yes | M |
| BLS_MAR_BMN_39_0795 | Tissue | 43.701 | 29.436 | BMN | MAR | no | yes | MC |
| BLS_MAR_BMN_39_0796 | Tissue | 43.701 | 29.436 | BMN | MAR | no | yes | M |
| BLS_MAR_BMN_39_0797 | Tissue | 43.701 | 29.436 | BMN | MAR | no | yes | M |
| BLS_MAR_BMN_39_0798 | Tissue | 43.701 | 29.436 | BMN | MAR | no | yes | M |

| | | | | | | | | |
|---|---|---|---|---|---|---|---|---|
| BLS_EST_CMN_62_1186 | Tissue | 45.645 | 36.497 | CMN | COT | no | yes | C |
| BLS_EST_CMN_62_1456 | Tissue | 45.645 | 36.497 | CMN | COT | no | yes | MC |
| BLS_EST_CMN_62_1182 | Tissue | 45.645 | 36.497 | CMN | COT | no | yes | MC |
| BLS_EST_CMN_62_1183 | Tissue | 45.645 | 36.497 | CMN | COT | no | yes | MC |
| BLS_EST_CMN_62_1184 | Tissue | 45.645 | 36.497 | CMN | COT | no | yes | C |
| BLS_EST_CMN_62_1185 | Tissue | 45.645 | 36.497 | CMN | COT | no | yes | C |
| ATL_MAR_CNR_88_1475 | Tissue | 27.934 | -15.605 | CNR | MAR | no | yes | S |
| BAL_MAR_DKB_65_1486 | Tissue | 54.347 | 11.681 | DKB | MAR | no | yes | M |
| BAL_MAR_DKB_65_1487 | Tissue | 54.347 | 11.681 | DKB | MAR | no | yes | MC |
| ATL_MAR_FAD_02_0950 | Tissue | 43.562 | -1.517 | GAS | MAR | no | yes | M |
| ATL_MAR_FAD_02_0958 | Tissue | 43.562 | -1.517 | GAS | MAR | no | yes | M |
| ATL_MAR_FAD_02_0961 | Tissue | 43.562 | -1.517 | GAS | MAR | no | yes | M |
| ATL_MAR_FAD_02_0963 | Tissue | 43.562 | -1.517 | GAS | MAR | no | yes | M |
| ATL_MAR_FAD_02_0974 | Tissue | 43.562 | -1.517 | GAS | MAR | no | yes | MC |
| ATL_MAR_FAD_02_0979 | Tissue | 43.562 | -1.517 | GAS | MAR | no | yes | M |
| ATL_MAR_FAD_02_0989 | Tissue | 43.562 | -1.517 | GAS | MAR | no | yes | M |
| ATL_EST_FAD_01_0894 | Le Moan et al. (2016) | 43.514 | -1.494 | GAS | COT | no | yes | MCS |
| ATL_EST_FAD_01_0896 | Le Moan et al. (2016) | 43.514 | -1.494 | GAS | COT | no | yes | CS |
| ATL_EST_FAD_01_0897 | Le Moan et al. (2016) | 43.514 | -1.494 | GAS | COT | no | yes | MC |
| ATL_EST_FAD_01_0899 | Le Moan et al. (2016) | 43.514 | -1.494 | GAS | COT | no | yes | M |
| ATL_EST_FAD_01_0900 | Le Moan et al. (2016) | 43.514 | -1.494 | GAS | COT | no | yes | MCS |
| ATL_EST_FAD_01_0902 | Le Moan et al. (2016) | 43.514 | -1.494 | GAS | COT | no | yes | M |
| ATL_EST_FAD_01_0903 | Le Moan et al. (2016) | 43.514 | -1.494 | GAS | COT | no | yes | C |
| ATL_EST_FAD_01_0904 | Le Moan et al. (2016) | 43.514 | -1.494 | GAS | COT | no | yes | M |
| ATL_EST_FAD_01_0905 | Le Moan et al. (2016) | 43.514 | -1.494 | GAS | COT | no | yes | MC |
| ATL_EST_FAD_01_0907 | Le Moan et al. (2016) | 43.514 | -1.494 | GAS | COT | no | yes | MCS |
| ATL_EST_FAD_01_0910 | Le Moan et al. (2016) | 43.514 | -1.494 | GAS | COT | no | yes | MCS |
| ATL_EST_FAD_01_0912 | Le Moan et al. | 43.514 | -1.494 | GAS | COT | no | yes | CS |

| | | | | | | | | |
|---|---|---|---|---|---|---|---|---|
| | (2016) | | | | | | | |
| ATL_EST_FAD_01_0913 | Le Moan et al. (2016) | 43.514 | -1.494 | GAS | COT | no | yes | C |
| ATL_EST_FAD_01_0914 | Le Moan et al. (2016) | 43.514 | -1.494 | GAS | COT | no | yes | M |
| ATL_EST_FAD_01_0916 | Le Moan et al. (2016) | 43.514 | -1.494 | GAS | COT | no | yes | MC |
| ATL_EST_FAD_01_0920 | Le Moan et al. (2016) | 43.514 | -1.494 | GAS | COT | no | yes | M |
| ATL_EST_FAD_01_0922 | Le Moan et al. (2016) | 43.514 | -1.494 | GAS | COT | no | yes | C |
| ATL_EST_FAD_01_0923 | Le Moan et al. (2016) | 43.514 | -1.494 | GAS | COT | no | yes | M |
| ATL_EST_FAD_01_0924 | Le Moan et al. (2016) | 43.514 | -1.494 | GAS | COT | no | yes | MC |
| ATL_EST_FAD_01_0925 | Le Moan et al. (2016) | 43.514 | -1.494 | GAS | COT | no | yes | M |
| ATL_EST_FAD_01_0926 | Le Moan et al. (2016) | 43.514 | -1.494 | GAS | COT | no | yes | M |
| ATL_EST_FAD_01_0929 | Le Moan et al. (2016) | 43.514 | -1.494 | GAS | COT | no | yes | M |
| ATL_EST_FAD_01_0930 | Le Moan et al. (2016) | 43.514 | -1.494 | GAS | COT | no | yes | MC |
| ATL_EST_FAD_01_0931 | Le Moan et al. (2016) | 43.514 | -1.494 | GAS | COT | no | yes | M |
| ATL_EST_FAD_01_0932 | Le Moan et al. (2016) | 43.514 | -1.494 | GAS | COT | no | yes | M |
| ATL_EST_FAD_01_0934 | Le Moan et al. (2016) | 43.514 | -1.494 | GAS | COT | no | yes | MC |
| ATL_EST_FAD_01_0935 | Le Moan et al. (2016) | 43.514 | -1.494 | GAS | COT | no | yes | MCS |
| ATL_EST_FAD_01_0937 | Le Moan et al. (2016) | 43.514 | -1.494 | GAS | COT | no | yes | MC |
| ATL_EST_FAD_01_0938 | Le Moan et al. (2016) | 43.514 | -1.494 | GAS | COT | no | yes | M |
| ATL_EST_FAD_01_0939 | Le Moan et al. (2016) | 43.514 | -1.494 | GAS | COT | no | yes | MC |
| ATL_EST_FAD_66_1139 | Le Moan et al. (2016) | 43.526 | -1.506 | GAS | COT | no | yes | MC |

| | | | | | | | | |
|---|---|---|---|---|---|---|---|---|
| ATL_EST_FAD_66_1145 | Le Moan et al. (2016) | 43.526 | -1.506 | GAS | COT | no | yes | MCS |
| ATL_EST_FAD_66_1147 | Le Moan et al. (2016) | 43.526 | -1.506 | GAS | COT | no | yes | CS |
| ATL_EST_FAD_66_1148 | Le Moan et al. (2016) | 43.526 | -1.506 | GAS | COT | no | yes | M |
| ATL_EST_FAD_66_1149 | Le Moan et al. (2016) | 43.526 | -1.506 | GAS | COT | no | yes | M |
| ATL_EST_FAD_66_1150 | Le Moan et al. (2016) | 43.526 | -1.506 | GAS | COT | no | yes | M |
| ATL_EST_FAD_66_1151 | Le Moan et al. (2016) | 43.526 | -1.506 | GAS | COT | no | yes | CS |
| ATL_EST_FAD_66_1153 | Le Moan et al. (2016) | 43.526 | -1.506 | GAS | COT | no | yes | MCS |
| ATL_MAR_GAS_90_1511 | Le Moan et al. (2016) | 44.710 | -1.480 | GAS | MAR | no | yes | MC |
| ATL_MAR_GAS_90_1512 | Le Moan et al. (2016) | 44.710 | -1.480 | GAS | MAR | no | yes | M |
| ATL_MAR_GAS_90_1513 | Le Moan et al. (2016) | 44.710 | -1.480 | GAS | MAR | no | yes | M |
| ATL_MAR_GAS_90_1514 | Le Moan et al. (2016) | 44.710 | -1.480 | GAS | MAR | no | yes | MCS |
| ATL_MAR_GAS_90_1515 | Le Moan et al. (2016) | 44.710 | -1.480 | GAS | MAR | no | yes | M |
| ATL_MAR_GAS_90_1516 | Le Moan et al. (2016) | 44.710 | -1.480 | GAS | MAR | no | yes | M |
| ATL_MAR_GAS_90_1517 | Le Moan et al. (2016) | 44.710 | -1.480 | GAS | MAR | no | yes | M |
| ATL_MAR_GAS_90_1518 | Le Moan et al. (2016) | 44.710 | -1.480 | GAS | MAR | no | yes | M |
| ATL_MAR_GAS_18_0621 | Le Moan et al. (2016) | 45.507 | -2.869 | GAS | MAR | no | yes | M |
| ATL_MAR_GAS_18_0622 | Le Moan et al. (2016) | 45.507 | -2.869 | GAS | MAR | no | yes | M |
| ATL_MAR_GAS_18_0624 | Le Moan et al. (2016) | 45.507 | -2.869 | GAS | MAR | no | yes | M |
| ATL_MAR_GAS_18_0625 | Le Moan et al. (2016) | 45.507 | -2.869 | GAS | MAR | no | yes | M |
| ATL_MAR_GAS_18_0634 | Le Moan et al. | 45.507 | -2.869 | GAS | MAR | no | yes | M |

| | | | | | | | | |
|---|---|---|---|---|---|---|---|---|
| | (2016) | | | | | | | |
| ATL_MAR_GAS_18_0644 | Le Moan et al. (2016) | 45.507 | -2.869 | GAS | MAR | no | yes | M |
| ATL_MAR_GAS_18_0645 | Le Moan et al. (2016) | 45.507 | -2.869 | GAS | MAR | no | yes | M |
| ATL_MAR_GAS_18_0646 | Le Moan et al. (2016) | 45.507 | -2.869 | GAS | MAR | no | yes | M |
| ATL_MAR_GAS_18_0647 | Le Moan et al. (2016) | 45.507 | -2.869 | GAS | MAR | no | yes | M |
| ATL_MAR_GAS_18_0648 | Le Moan et al. (2016) | 45.507 | -2.869 | GAS | MAR | no | yes | M |
| ATL_MAR_GAS_25_0853 | Le Moan et al. (2016) | 44.465 | -1.514 | GAS | MAR | no | yes | M |
| ATL_MAR_GAS_25_0856 | Le Moan et al. (2016) | 44.465 | -1.514 | GAS | MAR | no | yes | M |
| ATL_MAR_GAS_25_0857 | Le Moan et al. (2016) | 44.465 | -1.514 | GAS | MAR | no | yes | M |
| ATL_MAR_GAS_25_0858 | Le Moan et al. (2016) | 44.465 | -1.514 | GAS | MAR | no | yes | M |
| ATL_MAR_GAS_25_0864 | Le Moan et al. (2016) | 44.465 | -1.514 | GAS | MAR | no | yes | M |
| ATL_MAR_GAS_25_0865 | Le Moan et al. (2016) | 44.465 | -1.514 | GAS | MAR | no | yes | M |
| ATL_MAR_GAS_06_1103 | Tissue | 45.364 | -1.594 | GAS | MAR | no | yes | M |
| ATL_MAR_GAS_06_1098 | Tissue | 45.364 | -1.594 | GAS | MAR | no | yes | M |
| ATL_MAR_GAS_06_1099 | Tissue | 45.364 | -1.594 | GAS | MAR | no | yes | M |
| ATL_MAR_GAS_06_1100 | Tissue | 45.364 | -1.594 | GAS | MAR | no | yes | MC |
| ATL_MAR_GAS_06_1102 | Tissue | 45.364 | -1.594 | GAS | MAR | no | yes | M |
| ATL_MAR_GAS_70_1199 | Tissue | 45.400 | -1.380 | GAS | MAR | no | yes | M |
| ATL_MAR_GAS_70_1200 | Tissue | 45.400 | -1.380 | GAS | MAR | no | yes | M |
| ATL_MAR_GAS_70_1201 | Tissue | 45.400 | -1.380 | GAS | MAR | no | yes | M |
| ATL_MAR_GAS_70_1202 | Tissue | 45.400 | -1.380 | GAS | MAR | no | yes | M |
| ATL_MAR_GAS_70_1203 | Tissue | 45.400 | -1.380 | GAS | MAR | no | yes | M |
| ATL_MAR_GAS_70_1204 | Tissue | 45.400 | -1.380 | GAS | MAR | no | yes | M |
| ATL_MAR_GAS_70_1205 | Tissue | 45.400 | -1.380 | GAS | MAR | no | yes | M |
| ATL_MAR_GAS_70_1206 | Tissue | 45.400 | -1.380 | GAS | MAR | no | yes | M |
| ATL_MAR_FAD_02_0995 | Tissue | 43.562 | -1.517 | GAS | MAR | no | yes | MCS |
| ATL_MAR_FAD_02_0969 | Tissue | 43.562 | -1.517 | GAS | MAR | no | yes | M |
| ATL_MAR_FAD_02_0970 | Tissue | 43.562 | -1.517 | GAS | MAR | no | yes | M |

| | | | | | | | | |
|---|---|---|---|---|---|---|---|---|
| ATL_MAR_FAD_02_0977 | Tissue | 43.562 | -1.517 | GAS | MAR | no | yes | M |
| ATL_MAR_FAD_02_0978 | Tissue | 43.562 | -1.517 | GAS | MAR | no | yes | M |
| ATL_MAR_GAS_71_1207 | Tissue | 45.400 | -1.380 | GAS | MAR | no | yes | M |
| ATL_MAR_GAS_71_1208 | Tissue | 45.400 | -1.380 | GAS | MAR | no | yes | M |
| ATL_MAR_GAS_71_1209 | Tissue | 45.400 | -1.380 | GAS | MAR | no | yes | M |
| ATL_MAR_GAS_71_1210 | Tissue | 45.400 | -1.380 | GAS | MAR | no | yes | M |
| ATL_MAR_GAS_71_1211 | Tissue | 45.400 | -1.380 | GAS | MAR | no | yes | M |
| ATL_MAR_GAS_71_1212 | Tissue | 45.400 | -1.380 | GAS | MAR | no | yes | M |
| ATL_MAR_GAS_71_1213 | Tissue | 45.400 | -1.380 | GAS | MAR | no | yes | M |
| ATL_MAR_GAS_71_1214 | Tissue | 45.400 | -1.380 | GAS | MAR | no | yes | M |
| ATL_MAR_GAS_22_0712 | Tissue | 44.667 | -1.575 | GAS | MAR | no | yes | M |
| ATL_MAR_GAS_22_0717 | Tissue | 44.667 | -1.575 | GAS | MAR | no | yes | M |
| ATL_MAR_GAS_22_0733 | Tissue | 44.667 | -1.575 | GAS | MAR | no | yes | M |
| ATL_MAR_GAS_22_0734 | Tissue | 44.667 | -1.575 | GAS | MAR | no | yes | M |
| ATL_MAR_GAS_22_0743 | Tissue | 44.667 | -1.575 | GAS | MAR | no | yes | M |
| ATL_MAR_GAS_22_0750 | Tissue | 44.667 | -1.575 | GAS | MAR | no | yes | M |
| ATL_MAR_GAS_22_0754 | Tissue | 44.667 | -1.575 | GAS | MAR | no | yes | M |
| ATL_MAR_GAS_73_1254 | Tissue | 43.477 | -1.630 | GAS | MAR | no | yes | M |
| ATL_MAR_GAS_73_1255 | Tissue | 43.477 | -1.630 | GAS | MAR | no | yes | M |
| ATL_MAR_GAS_73_1256 | Tissue | 43.477 | -1.630 | GAS | MAR | no | yes | M |
| ATL_MAR_GAS_73_1257 | Tissue | 43.477 | -1.630 | GAS | MAR | no | yes | MC |
| ATL_MAR_GAS_73_1258 | Tissue | 43.477 | -1.630 | GAS | MAR | no | yes | MC |
| ATL_MAR_GAS_73_1260 | Tissue | 43.477 | -1.630 | GAS | MAR | no | yes | M |
| ATL_MAR_GAS_73_1261 | Tissue | 43.477 | -1.630 | GAS | MAR | no | yes | M |
| ATL_MAR_GAS_73_1263 | Tissue | 43.477 | -1.630 | GAS | MAR | no | yes | M |
| ATL_MAR_GAS_73_1246 | Tissue | 43.477 | -1.630 | GAS | MAR | no | yes | M |
| ATL_MAR_GAS_73_1265 | Tissue | 43.477 | -1.630 | GAS | MAR | no | yes | M |
| ATL_MAR_GAS_73_1266 | Tissue | 43.477 | -1.630 | GAS | MAR | no | yes | M |
| ATL_MAR_GAS_73_1264 | Tissue | 43.477 | -1.630 | GAS | MAR | no | yes | M |
| ATL_MAR_GAS_73_1247 | Tissue | 43.477 | -1.630 | GAS | MAR | no | yes | M |
| ATL_MAR_GAS_73_1248 | Tissue | 43.477 | -1.630 | GAS | MAR | no | yes | MC |
| ATL_MAR_GAS_73_1249 | Tissue | 43.477 | -1.630 | GAS | MAR | no | yes | M |
| ATL_MAR_GAS_73_1250 | Tissue | 43.477 | -1.630 | GAS | MAR | no | yes | M |
| ATL_MAR_GAS_73_1251 | Tissue | 43.477 | -1.630 | GAS | MAR | no | yes | M |
| ATL_MAR_GAS_73_1252 | Tissue | 43.477 | -1.630 | GAS | MAR | no | yes | M |
| ATL_MAR_GAS_73_1253 | Tissue | 43.477 | -1.630 | GAS | MAR | no | yes | M |
| ATL_MAR_GAS_74_1277 | Tissue | 43.471 | -1.658 | GAS | MAR | no | yes | M |
| ATL_MAR_GAS_74_1278 | Tissue | 43.471 | -1.658 | GAS | MAR | no | yes | M |
| ATL_MAR_GAS_74_1279 | Tissue | 43.471 | -1.658 | GAS | MAR | no | yes | M |

| | | | | | | | | |
|---|---|---|---|---|---|---|---|---|
| ATL_MAR_GAS_74_1280 | Tissue | 43.471 | -1.658 | GAS | MAR | no | yes | M |
| ATL_MAR_GAS_74_1281 | Tissue | 43.471 | -1.658 | GAS | MAR | no | yes | M |
| ATL_MAR_GAS_74_1282 | Tissue | 43.471 | -1.658 | GAS | MAR | no | yes | MC |
| ATL_MAR_GAS_74_1284 | Tissue | 43.471 | -1.658 | GAS | MAR | no | yes | M |
| ATL_MAR_GAS_74_1285 | Tissue | 43.471 | -1.658 | GAS | MAR | no | yes | MC |
| ATL_MAR_GAS_74_1267 | Tissue | 43.471 | -1.658 | GAS | MAR | no | yes | M |
| ATL_MAR_GAS_74_1286 | Tissue | 43.471 | -1.658 | GAS | MAR | no | yes | M |
| ATL_MAR_GAS_74_1287 | Tissue | 43.471 | -1.658 | GAS | MAR | no | yes | M |
| ATL_MAR_GAS_74_1288 | Tissue | 43.471 | -1.658 | GAS | MAR | no | yes | MC |
| ATL_MAR_GAS_74_1291 | Tissue | 43.471 | -1.658 | GAS | MAR | no | yes | M |
| ATL_MAR_GAS_74_1292 | Tissue | 43.471 | -1.658 | GAS | MAR | no | yes | M |
| ATL_MAR_GAS_74_1294 | Tissue | 43.471 | -1.658 | GAS | MAR | no | yes | M |
| ATL_MAR_GAS_74_1268 | Tissue | 43.471 | -1.658 | GAS | MAR | no | yes | M |
| ATL_MAR_GAS_74_1296 | Tissue | 43.471 | -1.658 | GAS | MAR | no | yes | MC |
| ATL_MAR_GAS_74_1298 | Tissue | 43.471 | -1.658 | GAS | MAR | no | yes | M |
| ATL_MAR_GAS_74_1299 | Tissue | 43.471 | -1.658 | GAS | MAR | no | yes | M |
| ATL_MAR_GAS_74_1300 | Tissue | 43.471 | -1.658 | GAS | MAR | no | yes | M |
| ATL_MAR_GAS_74_1301 | Tissue | 43.471 | -1.658 | GAS | MAR | no | yes | MC |
| ATL_MAR_GAS_74_1302 | Tissue | 43.471 | -1.658 | GAS | MAR | no | yes | M |
| ATL_MAR_GAS_74_1304 | Tissue | 43.471 | -1.658 | GAS | MAR | no | yes | M |
| ATL_MAR_GAS_74_1305 | Tissue | 43.471 | -1.658 | GAS | MAR | no | yes | M |
| ATL_MAR_GAS_74_1269 | Tissue | 43.471 | -1.658 | GAS | MAR | no | yes | M |
| ATL_MAR_GAS_74_1306 | Tissue | 43.471 | -1.658 | GAS | MAR | no | yes | M |
| ATL_MAR_GAS_74_1308 | Tissue | 43.471 | -1.658 | GAS | MAR | no | yes | M |
| ATL_MAR_GAS_74_1309 | Tissue | 43.471 | -1.658 | GAS | MAR | no | yes | M |
| ATL_MAR_GAS_74_1310 | Tissue | 43.471 | -1.658 | GAS | MAR | no | yes | M |
| ATL_MAR_GAS_74_1311 | Tissue | 43.471 | -1.658 | GAS | MAR | no | yes | M |
| ATL_MAR_GAS_74_1312 | Tissue | 43.471 | -1.658 | GAS | MAR | no | yes | M |
| ATL_MAR_GAS_74_1313 | Tissue | 43.471 | -1.658 | GAS | MAR | no | yes | MC |
| ATL_MAR_GAS_74_1314 | Tissue | 43.471 | -1.658 | GAS | MAR | no | yes | M |
| ATL_MAR_GAS_74_1315 | Tissue | 43.471 | -1.658 | GAS | MAR | no | yes | M |
| ATL_MAR_GAS_74_1270 | Tissue | 43.471 | -1.658 | GAS | MAR | no | yes | M |
| ATL_MAR_GAS_74_1316 | Tissue | 43.471 | -1.658 | GAS | MAR | no | yes | M |
| ATL_MAR_GAS_74_1271 | Tissue | 43.471 | -1.658 | GAS | MAR | no | yes | M |
| ATL_MAR_GAS_74_1272 | Tissue | 43.471 | -1.658 | GAS | MAR | no | yes | M |
| ATL_MAR_GAS_74_1273 | Tissue | 43.471 | -1.658 | GAS | MAR | no | yes | M |
| ATL_MAR_GAS_74_1274 | Tissue | 43.471 | -1.658 | GAS | MAR | no | yes | MC |
| ATL_MAR_GAS_74_1275 | Tissue | 43.471 | -1.658 | GAS | MAR | no | yes | M |
| ATL_MAR_GAS_75_1327 | Tissue | 43.467 | -1.671 | GAS | MAR | no | yes | M |

| | | | | | | | | |
|---|---|---|---|---|---|---|---|---|
| ATL_MAR_GAS_75_1333 | Tissue | 43.467 | -1.671 | GAS | MAR | no | yes | M |
| ATL_MAR_GAS_75_1338 | Tissue | 43.467 | -1.671 | GAS | MAR | no | yes | M |
| ATL_MAR_GAS_75_1340 | Tissue | 43.467 | -1.671 | GAS | MAR | no | yes | M |
| ATL_MAR_GAS_76_1359 | Tissue | 43.466 | -1.674 | GAS | MAR | no | yes | M |
| ATL_MAR_GAS_76_1360 | Tissue | 43.466 | -1.674 | GAS | MAR | no | yes | MC |
| ATL_MAR_GAS_76_1361 | Tissue | 43.466 | -1.674 | GAS | MAR | no | yes | M |
| ATL_MAR_GAS_76_1362 | Tissue | 43.466 | -1.674 | GAS | MAR | no | yes | M |
| ATL_MAR_GAS_76_1363 | Tissue | 43.466 | -1.674 | GAS | MAR | no | yes | M |
| ATL_MAR_GAS_76_1364 | Tissue | 43.466 | -1.674 | GAS | MAR | no | yes | M |
| ATL_MAR_GAS_76_1365 | Tissue | 43.466 | -1.674 | GAS | MAR | no | yes | M |
| ATL_MAR_GAS_76_1366 | Tissue | 43.466 | -1.674 | GAS | MAR | no | yes | M |
| ATL_MAR_GAS_76_1367 | Tissue | 43.466 | -1.674 | GAS | MAR | no | yes | M |
| ATL_MAR_GAS_76_1369 | Tissue | 43.466 | -1.674 | GAS | MAR | no | yes | M |
| ATL_MAR_GAS_76_1374 | Tissue | 43.466 | -1.674 | GAS | MAR | no | yes | M |
| ATL_MAR_GAS_76_1377 | Tissue | 43.466 | -1.674 | GAS | MAR | no | yes | M |
| ATL_MAR_GAS_76_1350 | Tissue | 43.466 | -1.674 | GAS | MAR | no | yes | M |
| ATL_MAR_GAS_77_1399 | Tissue | 43.484 | -1.650 | GAS | MAR | no | yes | M |
| ATL_MAR_GAS_76_1378 | Tissue | 43.466 | -1.674 | GAS | MAR | no | yes | M |
| ATL_MAR_GAS_76_1381 | Tissue | 43.466 | -1.674 | GAS | MAR | no | yes | M |
| ATL_MAR_GAS_76_1382 | Tissue | 43.466 | -1.674 | GAS | MAR | no | yes | M |
| ATL_MAR_GAS_76_1351 | Tissue | 43.466 | -1.674 | GAS | MAR | no | yes | M |
| ATL_MAR_GAS_76_1352 | Tissue | 43.466 | -1.674 | GAS | MAR | no | yes | M |
| ATL_MAR_GAS_76_1385 | Tissue | 43.466 | -1.674 | GAS | MAR | no | yes | M |
| ATL_MAR_GAS_76_1389 | Tissue | 43.466 | -1.674 | GAS | MAR | no | yes | M |
| ATL_MAR_GAS_76_1354 | Tissue | 43.466 | -1.674 | GAS | MAR | no | yes | M |
| ATL_MAR_GAS_76_1356 | Tissue | 43.466 | -1.674 | GAS | MAR | no | yes | M |
| ATL_MAR_GAS_77_1407 | Tissue | 43.484 | -1.650 | GAS | MAR | no | yes | M |
| ATL_MAR_GAS_77_1392 | Tissue | 43.484 | -1.650 | GAS | MAR | no | yes | M |
| ATL_MAR_GAS_77_1408 | Tissue | 43.484 | -1.650 | GAS | MAR | no | yes | M |
| ATL_MAR_GAS_77_1415 | Tissue | 43.484 | -1.650 | GAS | MAR | no | yes | MC |
| ATL_MAR_GAS_77_1419 | Tissue | 43.484 | -1.650 | GAS | MAR | no | yes | M |
| ATL_MAR_GAS_77_1397 | Tissue | 43.484 | -1.650 | GAS | MAR | no | yes | M |
| ATL_EST_FAD_78_1421 | Tissue | 43.525 | -1.506 | GAS | COT | no | yes | M |
| ATL_EST_FAD_78_1423 | Tissue | 43.525 | -1.506 | GAS | COT | no | yes | M |
| ATL_EST_FAD_78_1424 | Tissue | 43.525 | -1.506 | GAS | COT | no | yes | M |
| ATL_MAR_GAS_25_0821 | Tissue | 44.465 | -1.514 | GAS | MAR | no | yes | M |
| ATL_MAR_GAS_25_0830 | Tissue | 44.465 | -1.514 | GAS | MAR | no | yes | M |
| ATL_MAR_GAS_25_0835 | Tissue | 44.465 | -1.514 | GAS | MAR | no | yes | M |
| ATL_MAR_GAS_25_0838 | Tissue | 44.465 | -1.514 | GAS | MAR | no | yes | M |

| | | | | | | | | |
|---|---|---|---|---|---|---|---|---|
| ATL_MAR_GAS_25_0840 | Tissue | 44.465 | -1.514 | GAS | MAR | no | yes | M |
| ATL_MAR_GAS_25_0866 | Tissue | 44.465 | -1.514 | GAS | MAR | no | yes | M |
| ATL_MAR_FAD_02_0953 | Tissue | 43.562 | -1.517 | GAS | MAR | yes | yes | M |
| ATL_MAR_FAD_02_0957 | Tissue | 43.562 | -1.517 | GAS | MAR | yes | yes | M |
| ATL_MAR_FAD_02_0968 | Tissue | 43.562 | -1.517 | GAS | MAR | yes | yes | M |
| ATL_EST_FAD_01_0901 | Le Moan et al. (2016) | 43.514 | -1.494 | GAS | COT | yes | yes | C |
| ATL_EST_FAD_01_0911 | Le Moan et al. (2016) | 43.514 | -1.494 | GAS | COT | yes | yes | C |
| ATL_MAR_GAS_73_1259 | Tissue | 43.477 | -1.630 | GAS | COT | yes | yes | M |
| ATL_MAR_GAS_73_1262 | Tissue | 43.477 | -1.630 | GAS | COT | yes | yes | M |
| ATL_MAR_FAD_02_0947 | Tissue | 43.437 | -0.660 | GAS | COT | yes | yes | CS |
| ATL_MAR_FAD_02_0948 | Tissue | 43.437 | -0.660 | GAS | COT | yes | yes | C |
| ATL_MAR_FAD_02_0966 | Tissue | 43.437 | -0.660 | GAS | COT | yes | yes | CS |
| MED_LAG_GDL_82_1519 | Le Moan et al. (2016) | 43.576 | 4.018 | GDL | COT | no | yes | C |
| MED_LAG_GDL_82_1520 | Le Moan et al. (2016) | 43.576 | 4.018 | GDL | COT | no | yes | C |
| MED_LAG_GDL_82_1521 | Le Moan et al. (2016) | 43.576 | 4.018 | GDL | COT | no | yes | C |
| MED_LAG_GDL_82_1522 | Le Moan et al. (2016) | 43.576 | 4.018 | GDL | COT | no | yes | C |
| MED_LAG_GDL_82_1523 | Le Moan et al. (2016) | 43.576 | 4.018 | GDL | COT | no | yes | C |
| MED_LAG_GDL_82_1524 | Le Moan et al. (2016) | 43.576 | 4.018 | GDL | COT | no | yes | C |
| MED_LAG_GDL_82_1525 | Le Moan et al. (2016) | 43.576 | 4.018 | GDL | COT | no | yes | C |
| MED_LAG_GDL_82_1526 | Le Moan et al. (2016) | 43.576 | 4.018 | GDL | COT | no | yes | C |
| MED_LAG_GDL_82_1527 | Le Moan et al. (2016) | 43.576 | 4.018 | GDL | COT | no | yes | C |
| MED_LAG_GDL_82_1528 | Le Moan et al. (2016) | 43.576 | 4.018 | GDL | COT | no | yes | C |
| MED_LAG_GDL_82_1529 | Le Moan et al. (2016) | 43.576 | 4.018 | GDL | COT | no | yes | C |
| MED_LAG_GDL_82_1530 | Le Moan et al. (2016) | 43.576 | 4.018 | GDL | COT | no | yes | C |
| MED_LAG_GDL_82_1531 | Le Moan et al. | 43.576 | 4.018 | GDL | COT | no | yes | C |

| | | | | | | | | |
|---|---|---|---|---|---|---|---|---|
| | (2016) | | | | | | | |
| MED_LAG_GDL_82_1532 | Le Moan et al. (2016) | 43.576 | 4.018 | GDL | COT | no | yes | C |
| MED_LAG_GDL_82_1533 | Le Moan et al. (2016) | 43.576 | 4.018 | GDL | COT | no | yes | C |
| MED_LAG_GDL_82_1534 | Le Moan et al. (2016) | 43.576 | 4.018 | GDL | COT | no | yes | C |
| MED_LAG_GDL_82_1535 | Le Moan et al. (2016) | 43.576 | 4.018 | GDL | COT | no | yes | C |
| MED_LAG_GDL_82_1536 | Le Moan et al. (2016) | 43.576 | 4.018 | GDL | COT | no | yes | C |
| MED_LAG_GDL_82_1537 | Le Moan et al. (2016) | 43.576 | 4.018 | GDL | COT | no | yes | C |
| MED_LAG_GDL_82_1538 | Le Moan et al. (2016) | 43.576 | 4.018 | GDL | COT | no | yes | C |
| MED_LAG_GDL_83_1134 | Le Moan et al. (2016) | 43.576 | 4.018 | GDL | COT | no | yes | C |
| MED_LAG_GDL_83_1135 | Le Moan et al. (2016) | 43.576 | 4.018 | GDL | COT | no | yes | C |
| MED_LAG_GDL_83_1136 | Le Moan et al. (2016) | 43.576 | 4.018 | GDL | COT | no | yes | C |
| MED_LAG_GDL_83_1539 | Le Moan et al. (2016) | 43.576 | 4.018 | GDL | COT | no | yes | C |
| MED_LAG_GDL_82_1540 | Le Moan et al. (2016) | 43.576 | 4.018 | GDL | COT | no | yes | C |
| MED_LAG_GDL_82_1541 | Le Moan et al. (2016) | 43.576 | 4.018 | GDL | COT | no | yes | C |
| MED_LAG_GDL_82_1542 | Le Moan et al. (2016) | 43.576 | 4.018 | GDL | COT | no | yes | C |
| MED_LAG_GDL_82_1543 | Le Moan et al. (2016) | 43.576 | 4.018 | GDL | COT | no | yes | C |
| MED_LAG_GDL_82_1137 | Le Moan et al. (2016) | 43.576 | 4.018 | GDL | COT | no | yes | C |
| MED_LAG_GDL_82_1544 | Le Moan et al. (2016) | 43.576 | 4.018 | GDL | COT | no | yes | C |
| MED_LAG_GDL_82_1545 | Le Moan et al. (2016) | 43.576 | 4.018 | GDL | COT | no | yes | C |
| MED_LAG_GDL_82_1546 | Le Moan et al. (2016) | 43.576 | 4.018 | GDL | COT | no | yes | MC |

| | | | | | | | | |
|---|---|---|---|---|---|---|---|---|
| MED_MAR_GDL_81_1547 | Le Moan et al. (2016) | 43.199 | 3.613 | GDL | MAR | no | yes | MC |
| MED_MAR_GDL_81_1548 | Le Moan et al. (2016) | 43.199 | 3.613 | GDL | MAR | no | yes | M |
| MED_MAR_GDL_81_1124 | Le Moan et al. (2016) | 43.199 | 3.613 | GDL | MAR | no | yes | MC |
| MED_MAR_GDL_81_1549 | Le Moan et al. (2016) | 43.199 | 3.613 | GDL | MAR | no | yes | MC |
| MED_MAR_GDL_81_1550 | Le Moan et al. (2016) | 43.199 | 3.613 | GDL | MAR | no | yes | MC |
| MED_MAR_GDL_81_1551 | Le Moan et al. (2016) | 43.199 | 3.613 | GDL | MAR | no | yes | MC |
| MED_MAR_GDL_81_1125 | Le Moan et al. (2016) | 43.199 | 3.613 | GDL | MAR | no | yes | MC |
| MED_MAR_GDL_81_1127 | Le Moan et al. (2016) | 43.199 | 3.613 | GDL | MAR | no | yes | MC |
| MED_MAR_GDL_81_1128 | Le Moan et al. (2016) | 43.199 | 3.613 | GDL | MAR | no | yes | M |
| MED_MAR_GDL_81_1130 | Le Moan et al. (2016) | 43.199 | 3.613 | GDL | MAR | no | yes | M |
| MED_MAR_GDL_81_1131 | Le Moan et al. (2016) | 43.199 | 3.613 | GDL | MAR | no | yes | MC |
| MED_MAR_GDL_81_1133 | Le Moan et al. (2016) | 43.199 | 3.613 | GDL | MAR | no | yes | M |
| MED_MAR_GDL_81_0053 | Le Moan et al. (2016) | 43.329 | 3.820 | GDL | MAR | no | yes | MC |
| MED_MAR_GDL_81_0056 | Le Moan et al. (2016) | 43.329 | 3.820 | GDL | MAR | no | yes | M |
| MED_MAR_GDL_81_0057 | Le Moan et al. (2016) | 43.329 | 3.820 | GDL | MAR | no | yes | M |
| MED_MAR_GDL_81_0058 | Le Moan et al. (2016) | 43.329 | 3.820 | GDL | MAR | no | yes | MC |
| MED_MAR_GDL_81_0060 | Le Moan et al. (2016) | 43.329 | 3.820 | GDL | MAR | no | yes | M |
| MED_MAR_GDL_81_0061 | Le Moan et al. (2016) | 43.329 | 3.820 | GDL | MAR | no | yes | M |
| MED_MAR_GDL_81_0062 | Le Moan et al. (2016) | 43.329 | 3.820 | GDL | MAR | no | yes | M |
| MED_MAR_GDL_81_0063 | Le Moan et al. | 43.329 | 3.820 | GDL | MAR | no | yes | M |

| | | | | | | | | |
|---|---|---|---|---|---|---|---|---|
| | (2016) | | | | | | | |
| MED_MAR_GDL_81_0066 | Le Moan et al. (2016) | 43.329 | 3.820 | GDL | MAR | no | yes | MC |
| MED_MAR_GDL_81_0069 | Le Moan et al. (2016) | 43.329 | 3.820 | GDL | MAR | no | yes | M |
| MED_MAR_GDL_81_0071 | Le Moan et al. (2016) | 43.329 | 3.820 | GDL | MAR | no | yes | M |
| MED_MAR_GDL_81_0072 | Le Moan et al. (2016) | 43.329 | 3.820 | GDL | MAR | no | yes | M |
| MED_MAR_GDL_81_0074 | Le Moan et al. (2016) | 43.329 | 3.820 | GDL | MAR | no | yes | MCS |
| MED_MAR_GDL_81_0075 | Le Moan et al. (2016) | 43.329 | 3.820 | GDL | MAR | no | yes | MC |
| MED_MAR_GDL_81_0078 | Le Moan et al. (2016) | 43.329 | 3.820 | GDL | MAR | no | yes | M |
| MED_MAR_GDL_81_0079 | Le Moan et al. (2016) | 43.329 | 3.820 | GDL | MAR | no | yes | M |
| MED_MAR_GDL_81_1122 | Le Moan et al. (2016) | 43.199 | 3.613 | GDL | MAR | yes | yes | MC |
| MED_MAR_GDL_81_1129 | Le Moan et al. (2016) | 43.199 | 3.613 | GDL | MAR | yes | yes | M |
| MED_MAR_GDL_81_1132 | Le Moan et al. (2016) | 43.199 | 3.613 | GDL | MAR | yes | yes | M |
| Eencr_1L | Tissue | 43.528 | 3.886 | GDL | COT | yes | no | C |
| Eencr_2L | Tissue | 43.528 | 3.886 | GDL | COT | yes | no | C |
| EencrLi1 | Tissue | 43.199 | 3.613 | GDL | MAR | yes | no | MC |
| MED_MAR_GDL_81_1123 | Le Moan et al. (2016) | 43.199 | 3.613 | GDL | MAR | yes | yes | M |
| EencrLi4 | Tissue | 43.528 | 3.886 | GDL | COT | yes | no | C |
| EencrLi5 | Tissue | 43.528 | 3.886 | GDL | COT | yes | no | MCS |
| EencrLi6 | Tissue | 43.528 | 3.886 | GDL | COT | yes | no | C |
| ATL_MAR_MSA_57_1164 | Tissue | 21.998 | -16.965 | MA1 | MAR | no | yes | S |
| ATL_MAR_MSA_57_1165 | Tissue | 21.998 | -16.965 | MA1 | MAR | no | yes | S |
| ATL_MAR_MSA_57_1169 | Tissue | 21.998 | -16.965 | MA1 | MAR | no | yes | S |
| ATL_MAR_MSA_58_1425 | Tissue | 24.154 | -15.997 | MA2 | MAR | no | yes | S |
| ATL_MAR_MSA_58_1426 | Tissue | 24.154 | -15.997 | MA2 | MAR | no | yes | S |
| ATL_MAR_MSA_58_1427 | Tissue | 24.154 | -15.997 | MA2 | MAR | no | yes | S |
| ATL_MAR_MSA_58_1428 | Tissue | 24.154 | -15.997 | MA2 | MAR | no | yes | S |
| ATL_MAR_MSA_59_1445 | Tissue | 28.685 | -11.218 | MA3 | MAR | no | yes | SM |

| | | | | | | | | |
|---|---|---|---|---|---|---|---|---|
| ATL_MAR_MSA_59_1449 | Tissue | 28.685 | -11.218 | MA3 | MAR | no | yes | MCS |
| ATL_MAR_MSA_59_1450 | Tissue | 28.685 | -11.218 | MA3 | MAR | no | yes | MCS |
| ATL_MAR_MNA_56_1170 | Tissue | 33.678 | -7.698 | MA4 | MAR | no | yes | MCS |
| ATL_MAR_MNA_56_1171 | Tissue | 33.678 | -7.698 | MA4 | MAR | no | yes | CS |
| ATL_MAR_MNA_56_1172 | Tissue | 33.678 | -7.698 | MA4 | MAR | no | yes | MCS |
| ATL_MAR_MNA_56_1173 | Tissue | 33.678 | -7.698 | MA4 | MAR | no | yes | MCS |
| ATL_MAR_MDN_63_1215 | Tissue | 50.786 | 1.479 | MDN | MAR | no | yes | MC |
| ATL_MAR_MDN_63_1216 | Tissue | 50.786 | 1.479 | MDN | MAR | no | yes | MC |
| ATL_MAR_MDN_63_1217 | Tissue | 50.786 | 1.479 | MDN | MAR | no | yes | MC |
| ATL_MAR_MDN_63_1218 | Tissue | 50.786 | 1.479 | MDN | MAR | no | yes | MC |
| ATL_EST_MDN_91_1510 | Tissue | 51.500 | 0.640 | MDN | COT | no | yes | C |
| ATL_EST_MDN_91_1506 | Tissue | 51.500 | 0.640 | MDN | COT | no | yes | MC |
| ATL_EST_MDN_91_1507 | Tissue | 51.500 | 0.640 | MDN | COT | no | yes | MC |
| ATL_EST_MDN_91_1508 | Tissue | 51.500 | 0.640 | MDN | COT | no | yes | MC |
| ATL_EST_MDN_91_1509 | Tissue | 51.500 | 0.640 | MDN | COT | no | yes | C |
| ATL_EST_NOR_86_1459 | Tissue | 59.702 | 10.551 | NOR | COT | no | yes | MC |
| ATL_EST_NOR_86_1460 | Tissue | 59.702 | 10.551 | NOR | COT | no | yes | MC |
| BAL_MAR_PLN_85_1461 | Tissue | 55.238 | 18.058 | PLN | MAR | no | yes | MC |
| BAL_MAR_PLN_85_1462 | Tissue | 55.238 | 18.058 | PLN | MAR | no | yes | MC |
| BAL_MAR_PLN_85_1463 | Tissue | 55.238 | 18.058 | PLN | MAR | no | yes | MC |
| ATL_MAR_PRS_89_1470 | Tissue | 36.945 | -8.550 | PRS | MAR | no | yes | MCS |
| ATL_MAR_PRS_89_1471 | Tissue | 36.945 | -8.550 | PRS | MAR | yes | yes | MCS |
| ATL_MAR_PRS_89_1472 | Tissue | 36.945 | -8.550 | PRS | MAR | yes | yes | MCS |
| ATL_MAR_PRS_89_1473 | Tissue | 36.945 | -8.550 | PRS | MAR | yes | no | MCS |
| ATL_EST_PRS_84_1467 | Tissue | 37.028 | -7.812 | PRS | COT | yes | yes | CS |
| ATL_EST_PRS_84_1468 | Tissue | 37.028 | -7.812 | PRS | COT | yes | yes | MCS |
| ATL_EST_PRS_84_1469 | Tissue | 37.028 | -7.812 | PRS | COT | yes | yes | MCS |
| ATL_EST_PRS_72_1453 | Tissue | 37.029 | -8.003 | PRS | COT | yes | yes | C |
| ATL_EST_PRS_72_1454 | Tissue | 37.029 | -8.003 | PRS | COT | yes | yes | MCS |
| ATL_EST_PRS_72_1455 | Tissue | 37.029 | -8.003 | PRS | COT | yes | yes | MCS |
| EencrFa1 | Tissue | 36.945 | -8.550 | PRS | MAR | yes | no | MCS |
| EencrFa2 | Tissue | 36.945 | -8.550 | PRS | MAR | yes | no | CS |
| MED_LAG_SIC_68_1233 | Tissue | 38.269 | 15.637 | SIC | COT | no | yes | C |
| MED_LAG_SIC_68_1234 | Tissue | 38.269 | 15.637 | SIC | COT | no | yes | C |
| MED_LAG_SIC_68_1235 | Tissue | 38.269 | 15.637 | SIC | COT | no | yes | C |
| MED_LAG_SIC_68_1236 | Tissue | 38.269 | 15.637 | SIC | COT | no | yes | C |
| MED_MAR_SIC_67_1227 | Tissue | 38.148 | 15.595 | SIC | MAR | no | yes | M |
| MED_MAR_SIC_67_1228 | Tissue | 38.148 | 15.595 | SIC | MAR | no | yes | MC |
| MED_MAR_SIC_67_1229 | Tissue | 38.148 | 15.595 | SIC | MAR | no | yes | SM |

| | | | | | | | | |
|---|---|---|---|---|---|---|---|---|
| MED_MAR_SIC_67_1230 | Tissue | 38.148 | 15.595 | SIC | MAR | no | yes | M |
| EencrMu2 | Tissue | 37.970 | -0.682 | SPN | MAR | yes | no | MC |
| EencrMu3 | Tissue | 37.970 | -0.682 | SPN | MAR | yes | no | M |
| EencrMu4 | Tissue | 37.970 | -0.682 | SPN | MAR | yes | no | MCS |
| EencrMu5 | Tissue | 37.970 | -0.682 | SPN | MAR | yes | no | MCS |
| EencrMu6 | Tissue | 37.970 | -0.682 | SPN | MAR | yes | no | MCS |
| MED_MAR_TNO_40_0187 | Tissue | 37.067 | 9.000 | TNO | MAR | no | yes | M |
| MED_MAR_TNO_40_0196 | Tissue | 37.067 | 9.000 | TNO | MAR | no | yes | M |
| MED_MAR_TNO_40_0197 | Tissue | 37.067 | 9.000 | TNO | MAR | no | yes | M |
| MED_LAG_TNO_53_0388 | Tissue | 37.183 | 9.850 | TNO | COT | no | yes | C |
| MED_LAG_TNO_55_1075 | Tissue | 37.167 | 9.667 | TNO | COT | no | yes | C |
| MED_LAG_TNO_55_1076 | Tissue | 37.167 | 9.667 | TNO | COT | no | yes | C |
| MED_LAG_TNO_53_0405 | Tissue | 37.183 | 9.850 | TNO | COT | no | yes | C |
| MED_LAG_TNO_55_1079 | Tissue | 37.167 | 9.667 | TNO | COT | no | yes | C |
| MED_LAG_TNO_55_1081 | Tissue | 37.167 | 9.667 | TNO | COT | no | yes | C |
| ATL_MAR_ZDA_60_1154 | Tissue | -34.530 | 25.660 | ZDA | MAR | no | yes | S |
| ATL_MAR_ZDA_60_1156 | Tissue | -34.530 | 25.660 | ZDA | MAR | no | yes | S |
| ATL_MAR_ZDA_60_1157 | Tissue | -34.530 | 25.660 | ZDA | MAR | no | yes | S |
| ATL_MAR_ZDA_61_1160 | Tissue | -34.530 | 25.660 | ZDA | MAR | no | yes | S |
| ATL_MAR_ZDA_61_1161 | Tissue | -34.530 | 25.660 | ZDA | MAR | no | yes | S |
| ATL_MAR_ZDA_60_1155 | Tissue | -34.530 | 25.660 | ZDA | MAR | yes | yes | S |
| ATL_MAR_ZDA_61_1162 | Tissue | -34.530 | 25.660 | ZDA | MAR | yes | yes | S |
| ATL_MAR_ZDA_61_1159 | Tissue | -34.530 | 25.660 | ZDA | MAR | yes | yes | S |

**Supplementary Table S2: Location table.** Sampling locations used in the study. Samples were collected from both coastal and marine habitats in some locations.

| **Location** | **Description** |
|---|---|
| ALB | Morocco, Alboran Sea |
| BMN | Bulgaria, Black Sea |
| CMN | Kerch Strait, Black Sea |
| CNR | Canary islands, North-East Atlantic |
| DKB | Denmark, Baltic Sea |
| GAS | Bay of Biscay, North-East Atlantic |
| GDL | Gulf of Lion, Mediterranean Sea |
| MA1 | Southern Morocco, North-East Atlantic |
| MA2 | Southern Morocco, North-East Atlantic |

| | |
|---|---|
| MA3 | Nothern Morocco, North-East Atlantic |
| MA4 | Nothern Morocco, North-East Atlantic |
| MDN | English Channel, North-East Atlantic |
| NOR | Skagerrak, North-East Atlantic |
| PLN | Poland, Baltic Sea |
| PRS | Southern Portugal, North-East Atlantic |
| SIC | Sicily, Mediterranean Sea |
| SPN | Spain, Mediterranean Sea |
| TNO | Tunisia, Mediterranean Sea |
| ZDA | South Africa, South-East Atlantic |

**Supplementary Table S3: lineage differentiation based on SVs.** $F_{ST}$ between the three genetic clusters (*C*, *M* and S), calculated based on haplotype frequencies at SVs (*0* vs. *1* haplotypes).

| | SV region | | | | | | | | | | | | |
|---|---|---|---|---|---|---|---|---|---|---|---|---|---|
| **Genetic cluster comparison** | **CM068255** | **CM068256** | **CM068258** | **CM068260** | **CM068262** | **CM068265** | **CM068266** | **CM068267** | **CM068268** | **CM068270** | **CM068271** | **CM068273** | **CM068275** |
| **C vs M** | 0.707 | -0.006 | 0.845 | 0.025 | 0.059 | 0.452 | 0.015 | -0.005 | -0.005 | -0.004 | 0.016 | 0.031 | 0.037 |
| **C vs S** | -0.014 | 0.854 | -0.015 | 0.328 | 0.433 | 0.175 | 0.892 | 0.861 | 0.973 | 0.967 | 0.986 | 0.392 | 0.897 |
| **S vs M** | 0.68 | 0.933 | 0.818 | 0.485 | 0.588 | 0.681 | 0.971 | 0.866 | 0.959 | 0.983 | 0.933 | 0.488 | 0.727 |

#### Supplementary Figures

**Supplementary Fig. S1.** Genomic alignment dot plot showing the comparison between the chromosome-level assembly of *Engraulis encrasicolus* (top) and our genome assembly (right) subset to contain only scaffolds longer than 10 kb. Only shown here are alignment matches with similarity threshold of 50%. The base plot was generated using D-GENIES (https://github.com/genotoul-bioinfo/dgenies).

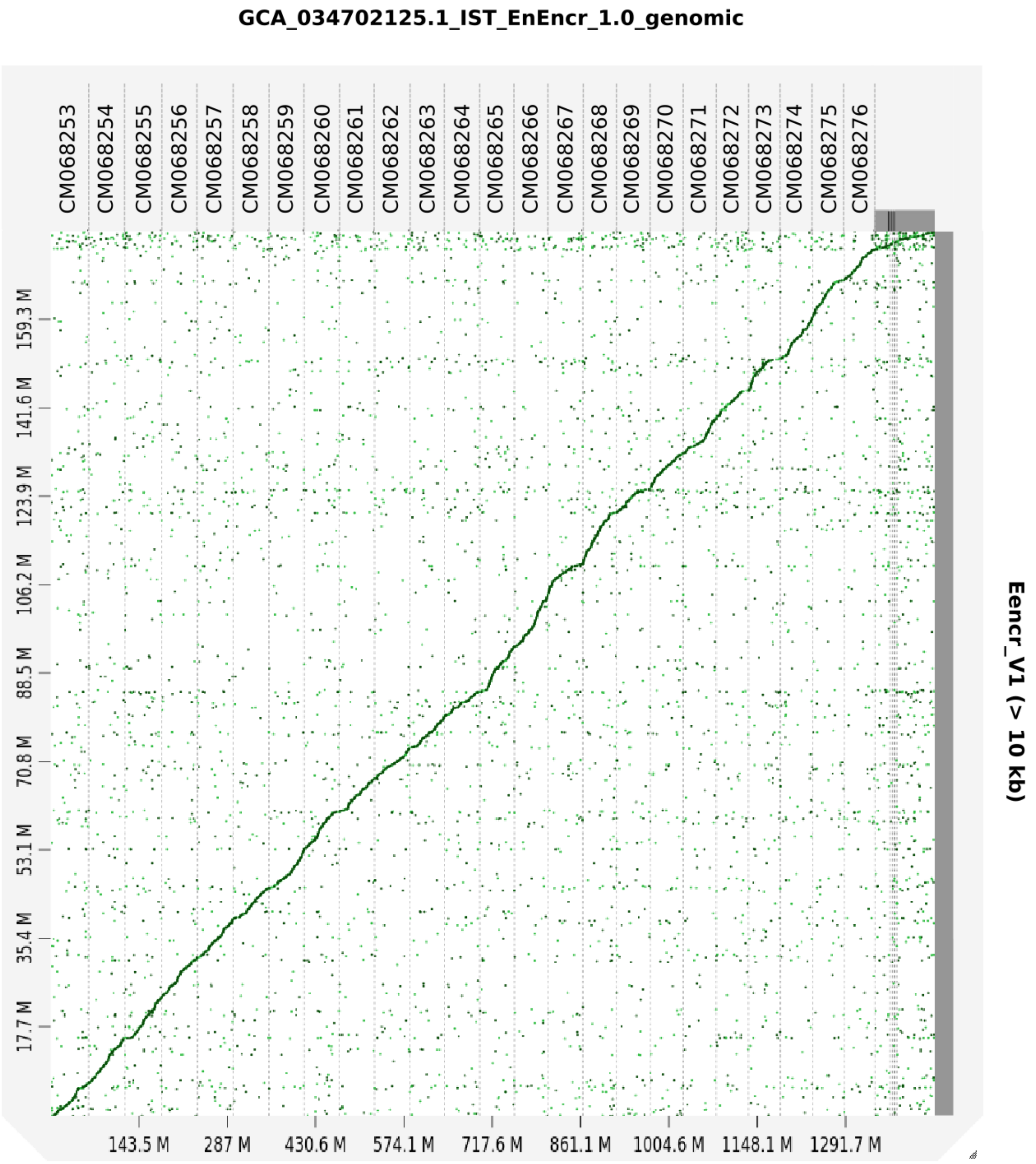

**Supplementary Fig. S2.** Chromosome-wide PCA conducted on all 24 chromosomes for whole-genome data (n=39). Square symbols represent samples from marine habitats, while diamond symbols represent samples from coastal habitats. Colours reflect individual admixture proportions based on genome-wide admixture analysis (**Fig 1B**).

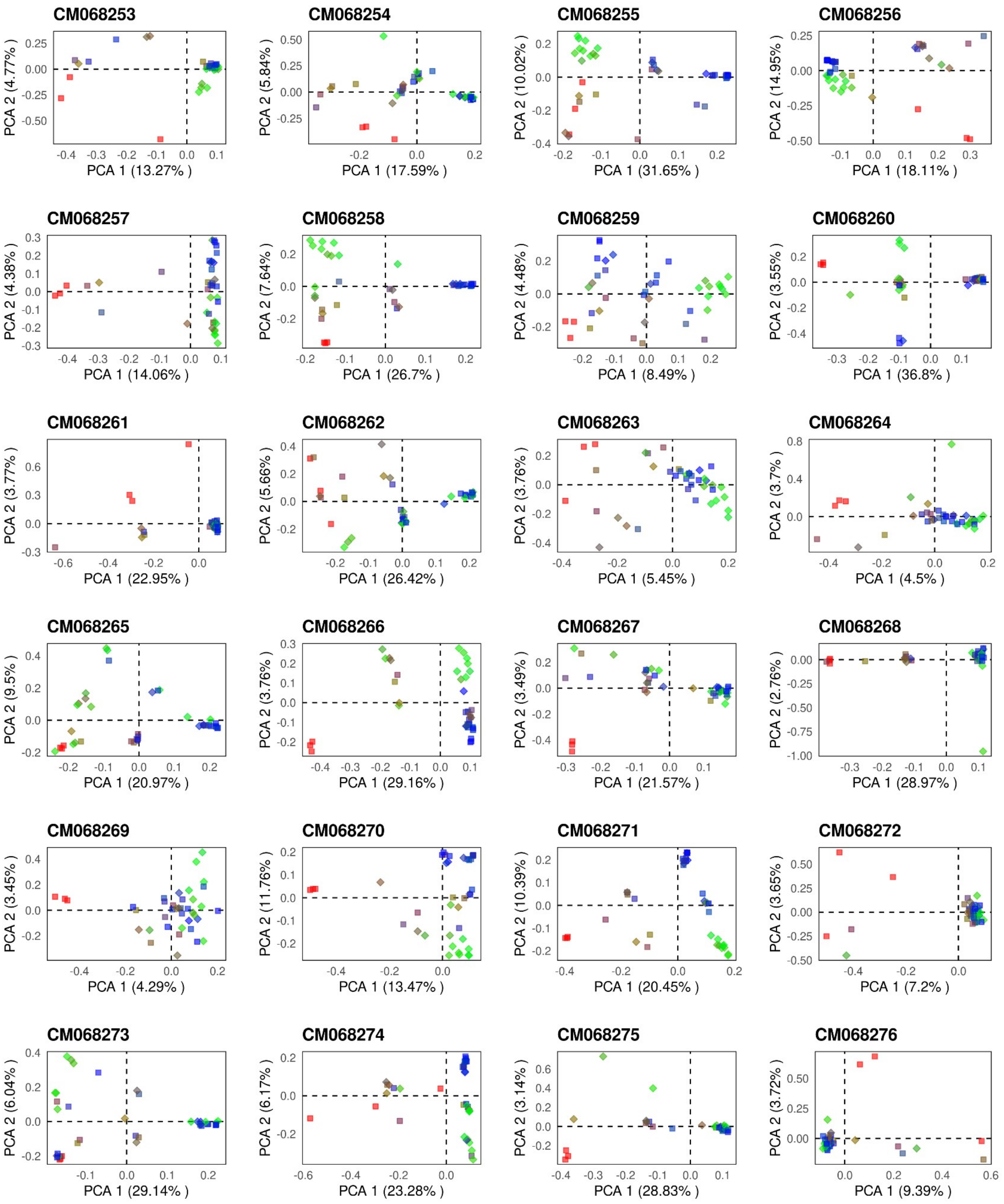

**Supplementary Fig. S3.** Assignment of SV genotypes for whole-genome data (n=39). Plots show genome-wide PCA conducted on 13 different chromosomes, where individuals were classified as *00* homokaryotes (pink), *01* heterokaryotes (salmon) or *11* homokaryotes (gold). Symbols represent habitat type (squares - marine, diamonds - coastal).

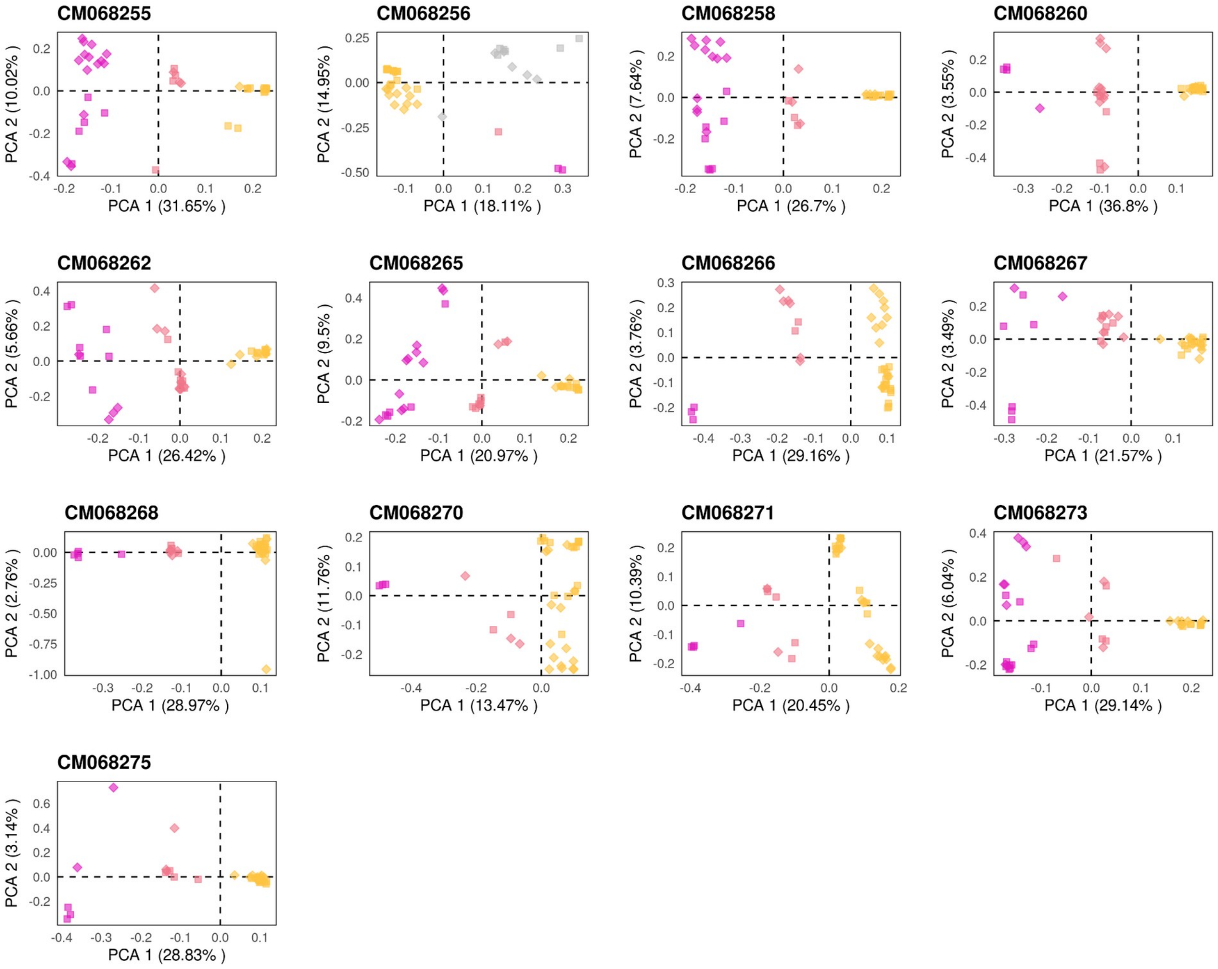

**Supplementary Fig. S4.** Chromosome-wide PCA conducted on all 24 chromosomes for RAD data (n=385). Square symbols represent samples from marine habitats, while diamond symbols represent samples from coastal habitats. Colours reflect individual admixture proportions based on genome-wide admixture analysis (**Fig 1B**).

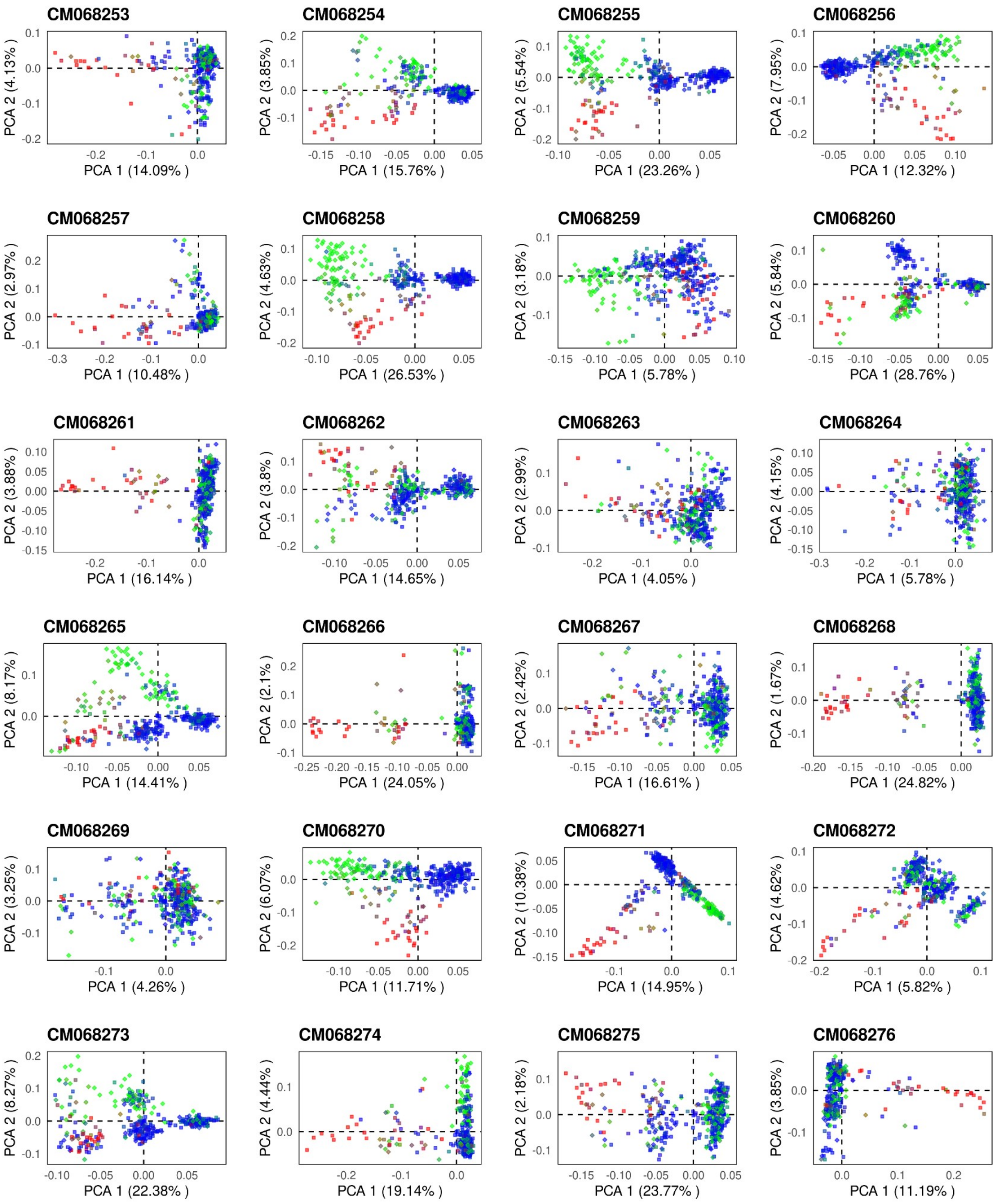

**Supplementary Fig. S5.** Assignment of SV genotypes for RAD data (n=385). Plots show genome-wide PCA conducted on 13 different chromosomes, where individuals were classified as *00* homokaryotes (pink), *01* heterokaryotes (salmon) or *11* homokaryotes (gold). Symbols represent habitat type (squares - marine, diamonds - coastal).

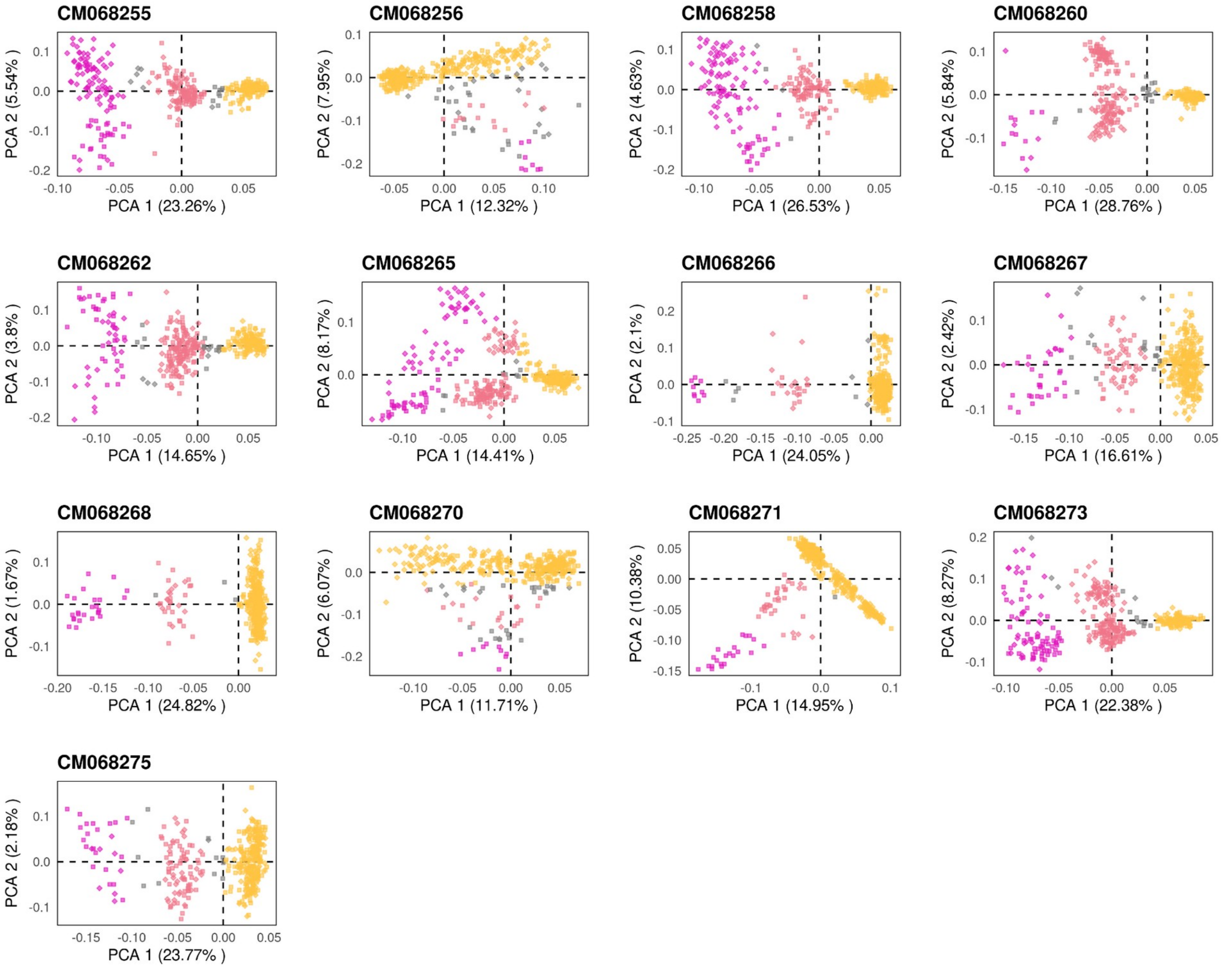

**Supplementary Fig. S6.** Genotypes at all SVs (rows) across all individuals (columns). Individuals are firstly grouped according to their locations, and secondly, according to their genome-wide ancestry categories. Due to space constraints, not all marine samples from the *GAS* location are shown. Colours indicate the genotype at a given SV and are the same colours as in **Supplementary Fig. S5**.

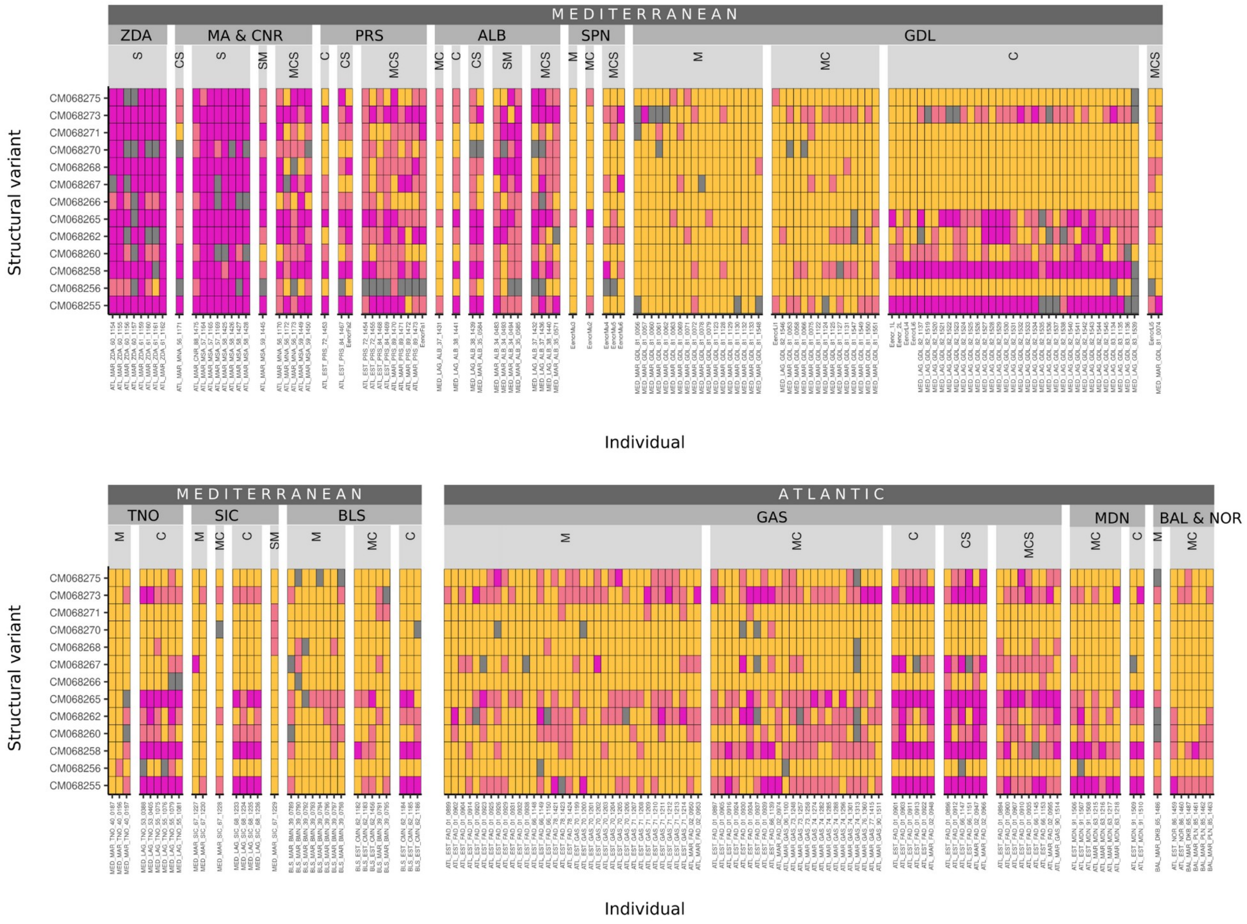

**Supplementary Fig. S7.** Neighbour-joining trees constructed in different genomic regions. For chromosomes carrying SVs, the region was limited to windows showing patterns of high LD (based on $F_{ST}$ and local PCA) and intermediate samples that present heterokaryotes are not displayed. For chromosomes without SVs, trees were constructed using SNPs on the entire chromosome. Leaf labels are coloured according to individual ancestry proportions and tip symbols (circles) correspond to SV genotype. Trees were plotted on the same vertical scale.

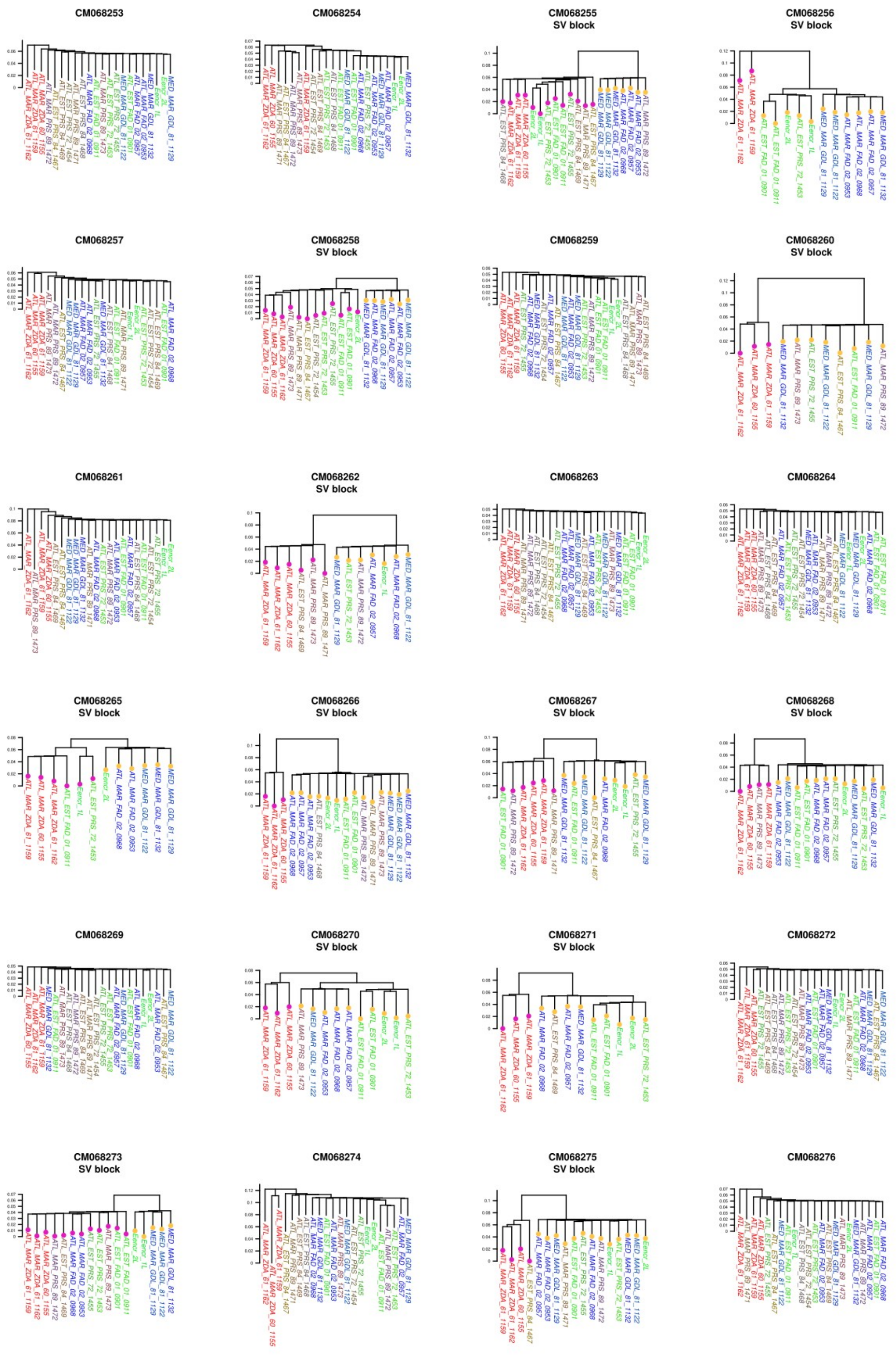

**Supplementary Fig. S8.** Haplotype frequencies for 13 SVs on different chromosomes (columns). Pie charts in (A) show frequencies for *0* haplotypes (pink) and *1* haplotypes (gold) in the marine cluster (blue background), coastal cluster (green background) and southern cluster (red background) (Mediterranean and Atlantic combined). Darker background colour in *M* and *C* indicates chromosomes that show elevated $F_{ST}$ when comparing marine and coastal individuals. Pie charts in (B) show sub-haplotype frequencies, with colours corresponding to *0* haplotypes (pink), *1a* haplotypes (brown) and *1b* haplotypes (gold). Grey sectors represent the proportion of samples that were not genotyped (NA).

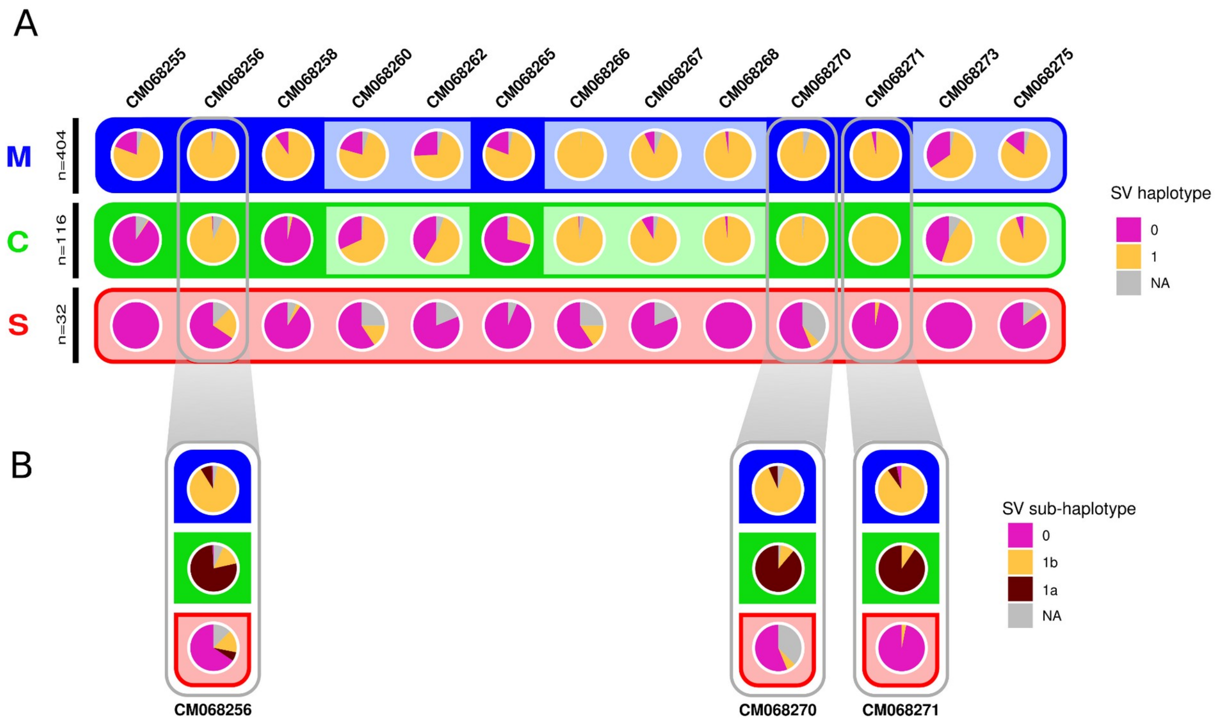

**Supplementary Fig. S9.** Distributions of genetic differentiation ($F_{ST}$) values in different regions of the genome (SVs or the background genome). The analysis compared two populations composed of three individuals each (from the same ancestry category).

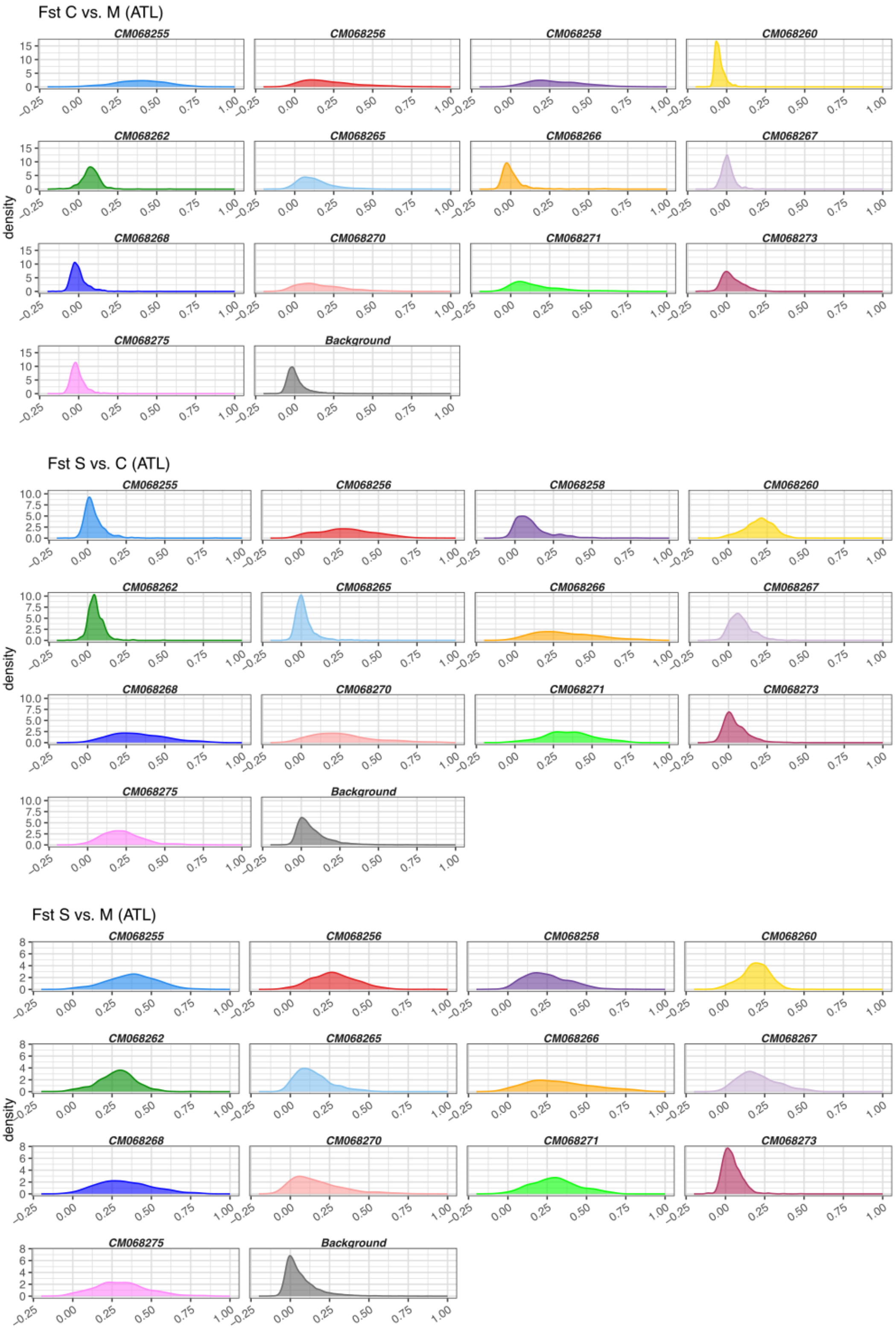

**Supplementary Fig. S10.** Distributions of absolute differentiation ($d_{XY}$) and nucleotide diversity ($\pi$) values in different SV regions. The analysis included three individuals that were assigned to the *00* genotype and three individuals that were assigned to the *11* genotype (except for the SV on CM068256 where only two *00* individuals were identified in the WGS dataset).

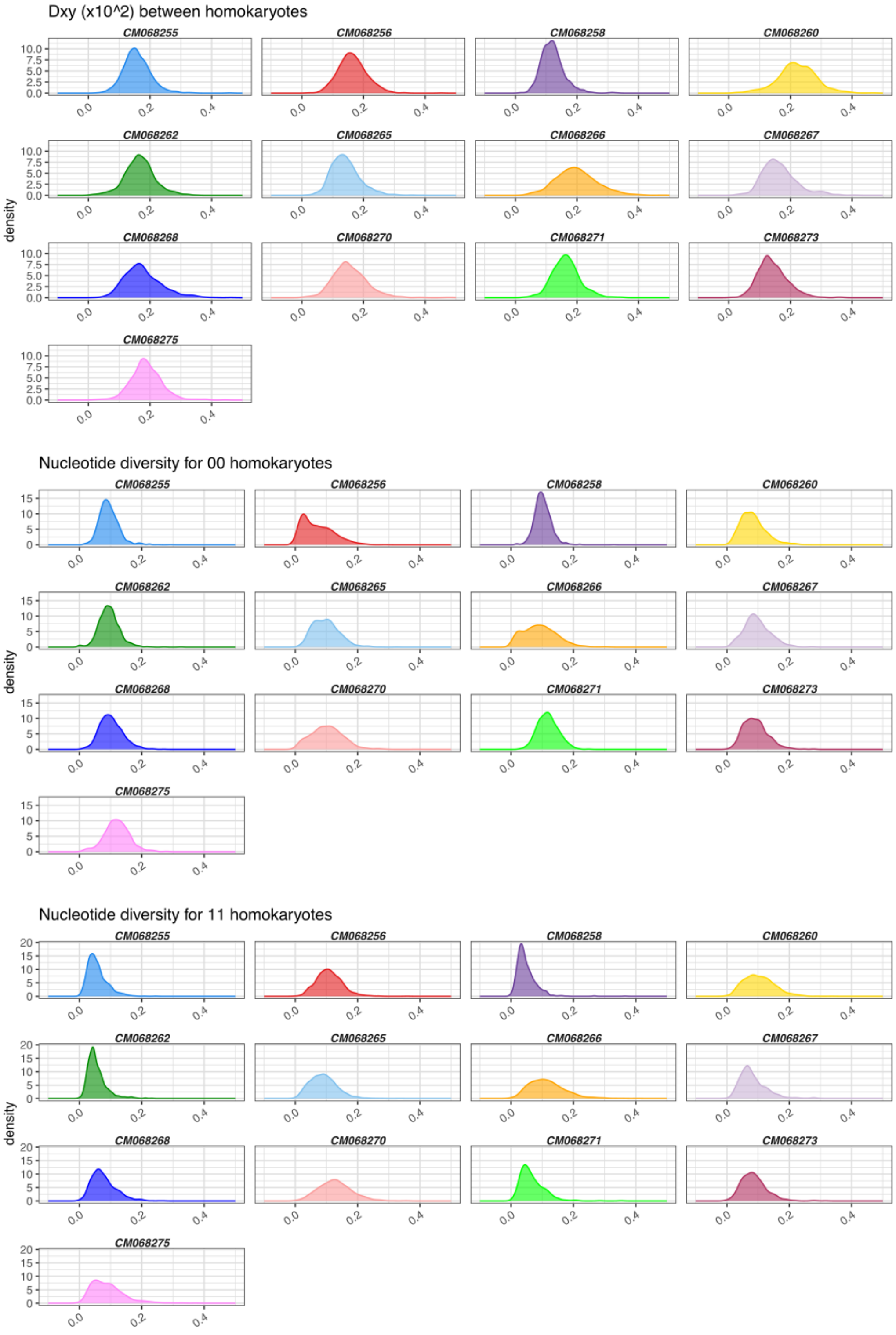

**Supplementary Fig. S11.** Distributions of observed heterozygosity (percentage of heterozygous sites per individual) in the 13 different SV regions. Points represent individuals from the WGS dataset, grouped according to their SV genotype (*00*, *01* or *11*).

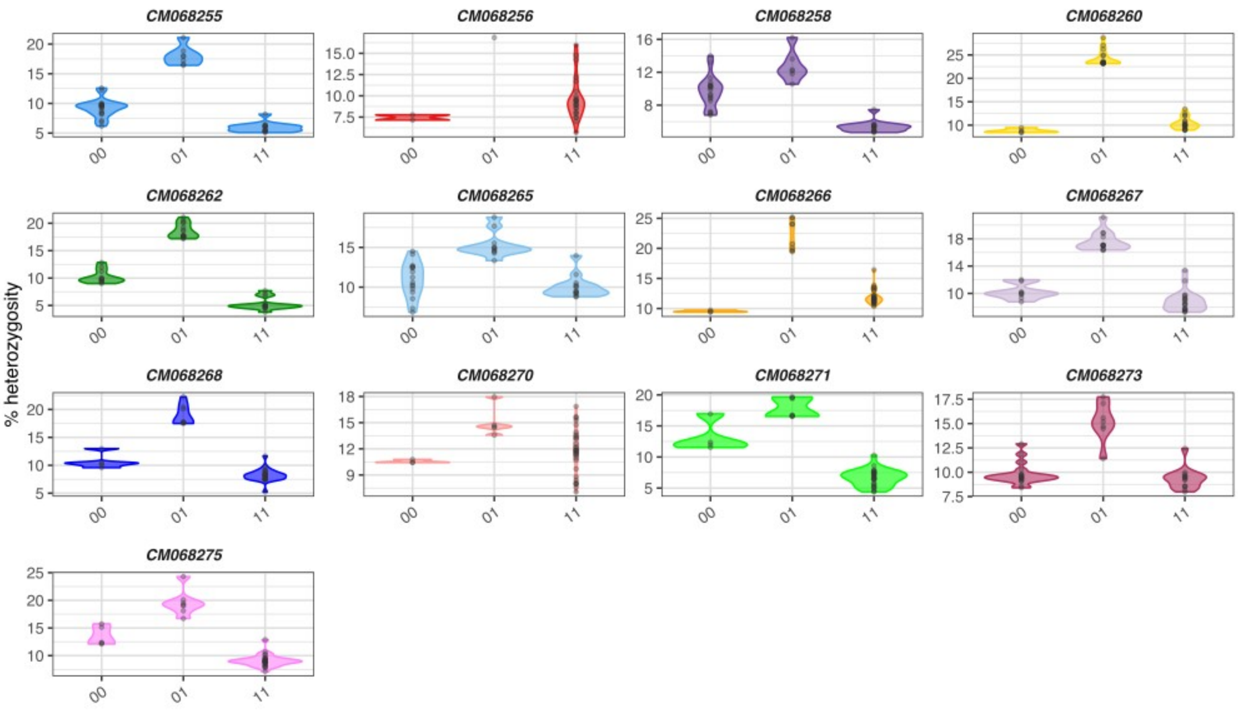

### Supplementary Appendix

Detailed results can be consulted in Anchovy_Results_Appendix.html. This report includes results about the alignment of RAD and WGS data, PCA conducted on different genomic regions, and the SVs that were genotyped.
